# HIV Vpr modulates the host DNA damage response at two independent steps to damage DNA and repress double-strand DNA break repair

**DOI:** 10.1101/2020.04.26.062349

**Authors:** Donna Li, Andrew Lopez, Carina Sandoval, Randilea Nichols Doyle, Oliver I Fregoso

## Abstract

The DNA damage response (DDR) is a signaling cascade that is vital to ensuring the fidelity of the host genome in the presence of genotoxic stress. Growing evidence has emphasized the importance of both activation and repression of the host DDR by diverse DNA and RNA viruses. Previous work has shown that HIV-1 is also capable of engaging the host DDR, primarily through the conserved accessory protein Vpr. However, the extent of this engagement has remained unclear. Here we show that HIV-1 and HIV-2 Vpr directly induce DNA damage and stall DNA replication, leading to the activation of several markers of double- and single-strand DNA breaks. Despite causing damage and activating the DDR, we found that Vpr repress the repair of double-strand breaks (DSB) by inhibiting homologous recombination (HR) and non-homologous end joining (NHEJ). Mutational analyses of Vpr revealed that DNA damage and DDR activation are independent from repression of HR and Vpr-mediated cell-cycle arrest. Moreover, we show that repression of HR does not require cell-cycle arrest but instead may precede this long-standing enigmatic Vpr phenotype. Together, our data uncover that Vpr globally modulates the host DDR at at least two independent steps; offering novel insight into the primary functions of lentiviral Vpr and the roles of the DNA damage response in lentiviral replication.

**IMPORTANCE:** The DNA damage response (DDR) is a signaling cascade that safeguards the genome from genotoxic agents, including human pathogens. However, the DDR has also been utilized by many pathogens, such as Human Immunodeficiency Virus (HIV), to enhance infection. To properly treat HIV positive individuals, we must understand how the virus usurps our own cellular processes. Here, we have found that an important yet poorly-understood gene in HIV, Vpr, targets the DDR at two unique steps: it causes damage and activates DDR signaling, and it represses the ability of cells to repair this damage, which we hypothesize is central to the primary function of Vpr. In clarifying these important functions of Vpr, our work highlights the multiple ways human pathogens engage the DDR, and further suggests that modulation of the DDR may be a novel way to help in the fight against HIV.

## INTRODUCTION

Primate lentiviruses encode accessory proteins that enhance viral (1). This is achieved through direct interactions with host proteins to usurp their cellular functions, or to antagonize their antiviral activity. HIV-1 encodes four accessory factors: Vpr, Vif, Vpu, and Nef. In addition, a subset of lentiviruses, including HIV-2, encode a paralog of Vpr called Vpx (2). Of all the lentiviral accessory genes, *vpr* is the only gene with a still unknown primary function.

Despite this, Vpr is critical for the infectivity of HIV and related primate lentiviruses. *In vivo*, viruses lacking Vpr are attenuated compared to wild type viruses, and the dominant viral species to emerge (i.e. most fit) have restored Vpr protein expression (3, 4). Furthermore, *vpr* is evolutionarily conserved by all extant primate lentiviruses (5). Together, this indicates that lentiviruses have maintained *vpr* for a highly important function. Of the many potential roles assigned to Vpr, activation of the host DNA damage response (DDR) and subsequent cell cycle arrest are the only phenotypes conserved by diverse Vpr orthologs (6–8). This conservation of function suggests that engagement of the DDR is central to Vpr function.

The DNA damage response (DDR) is a protein signaling cascade that ensures the fidelity of the genome. It consists of sensors, which recognize specific DNA lesions, mediators and transducers that transmit this signal of damaged DNA, and effectors, which directly execute a cellular response. Ataxia telangiectasia and Rad3 (ATR) (9), Ataxia telangiectasia mutated (ATM) (10), and DNA dependent protein kinase (DNA-PK) (11) are kinases at the head of the complex network that makes up the host DDR. The ATR kinase primarily responds to UV damage and replication stress, while ATM and DNA-PK participate in the repair of double-strand breaks (DSB) through homologous recombination (HR) and non-homologous end joining (NHEJ), respectively (12). However, due to the essential role of the DDR, a tremendous amount of crosstalk and redundancy exists between these kinases (13).

There is growing evidence that the DDR is important for viral replication, where it acts to both enhance and inhibit replication (14). For example, the DNA virus herpes simplex virus 1 (HSV-1) induces replication fork collapse at sites of oxidative damage (15). This leads to double-strand breaks (DSB), which initiate activation of the ATM repair pathway. HSV-1 infection also activates ATR, and the inactivation of either pathway severely compromises HSV-1 replication. RNA viruses also engage the DDR; for example, Rift Valley Fever Virus activates markers of DNA damage such as *γ*H2AX and upregulates the ATM pathway but represses the ATR pathway (16). Contrary to enhancing viral replication, DDR proteins, such as DNA-PK (17), can activate an antiviral state upon sensing cytoplasmic DNA, while etoposide-induced DNA damage stimulates interferon via STING, ATM, and NF-kB (18–22). Together, this highlights the potential roles for the DDR in innate antiviral immunity and in enhancing viral replication.

Vpr engages the DDR at multiple steps. First, it causes G2 cell cycle arrest both *in vivo* and *in vitro* (7, 23–26). This arrest is dependent on ATR signaling, as it is blocked by chemical inhibition of ATR (27). Moreover, Vpr-mediated cell cycle arrest requires interaction of Vpr with the Cul4A/DCAF1/DDB1 (CUL4A^DCAF1^) E3 ubiquitin ligase complex (28, 29), a cellular complex that is involved in many mechanisms of DNA repair (30, 31). Second, Vpr induces the expression, activation, and recruitment of DDR proteins, as assessed by immunofluorescence and western blot analysis (32–34). Finally, in addition to the CUL4A^DCAF1^ ubiquitin ligase complex, Vpr interacts with and degrades many host DDR proteins, including UNG2 (35, 36), HLTF (37, 38), SLX4 complex proteins MUS81 and EME1 (34, 39), EXO1 (40), TET2 (41), MCM10 (42), and SAMHD1 (5, 43). Yet despite being one of the most highly conserved and robust phenotypes associated with Vpr, how Vpr engages the DDR at so many levels remains unclear.

Using a combination of DNA damage response assays, we monitored the induction of DNA damage, the early signaling events following DDR activation, and the cellular consequences associated with DNA damage and DDR activation. We found that Vpr engages the DNA damage response at two independent steps: it causes DNA damage and activates DDR signaling, and it represses double-strand DNA break repair. Using a panel of HIV-1 and HIV-2 Vpr mutants, we were able to separate these Vpr functions to show that while Vpr-induced DNA damage is independent of most known Vpr-host protein interactions, repression of double-strand break repair is dependent on DCAF1 recruitment. Finally, we showed that repression of HR repair is not a consequence of Vpr-mediated G2 cell cycle arrest, but instead may be a driver of this long-standing Vpr phenotype. Our data indicate that lentiviruses both activate and repress the DDR via Vpr, and further characterize a novel phenotype of Vpr that can help explain many of the roles that have long been associated with Vpr.

## RESULTS

### HIV-1 and HIV-2 Vpr activate multiple DNA damage markers

The extent to which HIV Vpr engages the host DNA damage response (DDR) has not been critically examined. Therefore, we first asked if both HIV-1 and HIV-2 Vpr similarly activate the DDR. HIV-1 and HIV-2 are evolutionarily divergent primate lentiviruses that entered the human population through different non-human primate hosts (44). The Vpr proteins of these two viruses share only about 30-40% similarity, yet both cause cell cycle arrest (5, 7). Thus, if engagement of the DDR was central to the function of Vpr, we would expect that Vpr proteins from these two diverse human lentiviruses would also similarly activate the DDR. To test this, we delivered HIV-1 Q23-17 Vpr and HIV-2 Rod9 Vpr to U2OS cells via a recombinant adeno-associated virus (rAAV) vector system expressing 3X FLAG-tagged Vpr (6) and assayed for DDR markers twenty hours post infection by immunofluorescence (IF) for *γ*H2AX, a marker for DNA double- and single-strand breaks (DSB and SSB, respectively) (45), RPA32, a marker of SSB (46), and 53BP1, a late marker of DSB that is recruited to sites of damage by *γ*H2AX (47). In the presence of HIV-1 and HIV-2 Vpr, there were increased amounts of *γ*H2AX foci compared to the uninfected and empty vector controls (Fig. 1), which correlated with G2 arrest (Fig. S1A). Similar to *γ*H2AX, HIV-1 and HIV-2 Vpr expression also lead to increased levels of RPA32 and 53BP1 foci compared to uninfected and empty vector control cells, and produced fewer yet larger foci compared to the etoposide positive control, a topoisomerase II inhibitor (R) (Fig. 1). We also observed a distinct lack of colocalization between Vpr and markers of DNA damage (Fig. 1A), indicating that Vpr is not present at the potential sites of damage at this time point. Additionally, individual cells that expressed higher levels of Vpr did not have appreciably more DNA damage foci or mean fluorescence intensity (MFI) of *γ*H2AX, RPA32, or 53BP1 per cell when compared to cells with lower Vpr expression (Fig. 1A asterisks and Fig. S1B-E, respectively), suggesting that activation of these markers is saturated with low levels of Vpr. And while HIV-1 had significantly higher (P<0.03) levels of 53BP1 compared to HIV-2 Vpr, the levels of *γ*H2AX and RPA32 activation were the same for HIV-1 and HIV-2 as measured by MFI of individual cells (Fig. 1B).

**Figure 1.**
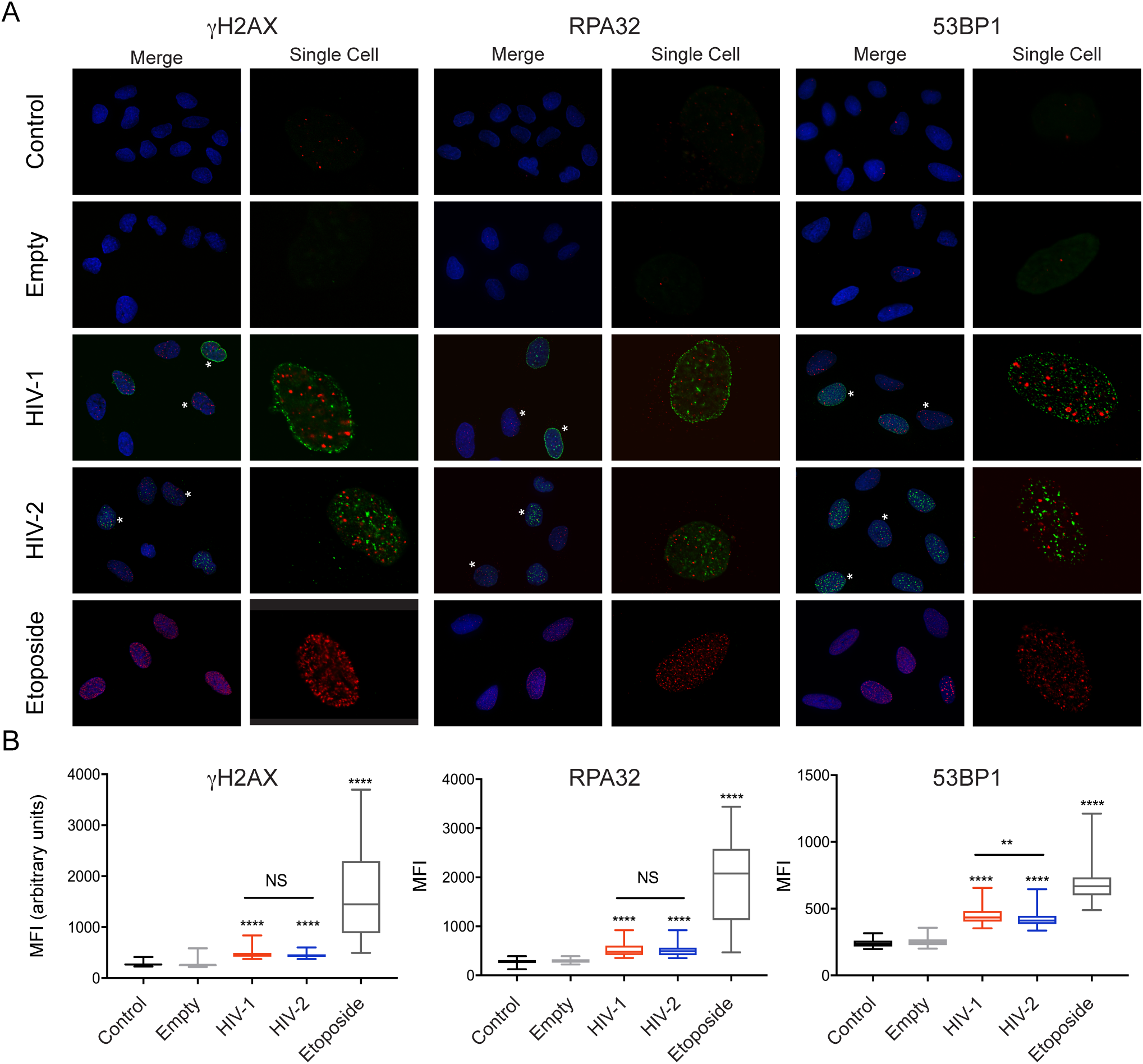
Activation of the DNA damage response is conserved between HIV-1 and HIV-2 Vpr. A. Representative immunofluorescence images of U2OS cells infected with rAAV expressing 3X FLAG-tagged HIV-1 and HIV-2 Vpr, control empty vector (no Vpr), or uninfected control for 20 hrs. Blue (DAPI) shows the nuclei, 3X-FLAG Vpr is shown in green, and the phosphorylated DNA damage markers (*γ*H2AX, RPA32, and 53BP1) are shown in red. Asterisks indicate cells with either high or low Vpr expression. The single cell images show only 3X-FLAG Vpr and corresponding DNA damage marker. Images taken at 63x magnification. B. Mean fluorescence intensity (MFI) of 100 cells per condition was quantified for all markers. Asterisks indicate statistical significance in comparison to empty vector control as determined by Kruskal-Wallis tests (NS, nonsignificant; *, *P*<0.05; **, *P*<0.01; ***, P<0.001; ****, *P*<0.0001; n=2, one representative experiment shown).

We also tested a number of HIV-1 and HIV-2 Vpr isolates to determine if activation of the DDR by HIV-1 and HIV-2 Vpr were isolate specific or conserved by the greater diversity of HIV Vpr proteins. These include representative Vpr isolates from HIV-1 groups M (subtype G), N, O, and P consensus sequences, as well as HIV-2 Vpr isolates from groups A, B, and divergent. We found that all HIV-1 and HIV-2 Vpr proteins tested caused cell cycle arrest and increased the number of *γ*H2AX foci, indicative of DDR activation (Fig. S2). In total, our data highlights that a conserved function of HIV-1 and HIV-2 Vpr is the activation of the same markers of single- and double-strand DNA damage.

### HIV-1 and HIV-2 Vpr expression damages DNA and induces replication stress

The formation of *γ*H2AX, RPA32, and 53BP1 foci in cells expressing HIV-1 and HIV-2 Vpr suggests the presence of both SSB and DSB. However, it is also possible that Vpr leads to activation of these markers without causing actual DNA damage. Previous studies to identify Vpr-induced DNA damage using pulse field gel electrophoresis, which only reveals DSB, have been contradictory (48, 49). Here we used the alkaline comet assay, which uses a high pH (>10) buffer to denature supercoiled DNA and single-cell gel electrophoresis to reveal damaged DNA fragments, including both SSB and DSB (50). U2OS cells were infected with rAAV-Vpr for twenty hours and the extent of DNA damage within individual cells was measured by calculating the percent tail DNA, which is proportionate to the amount of damaged DNA within a cell (Fig. 2A). While uninfected and empty vector control cells had little appreciable damage, both HIV-1 and HIV-2 Vpr expression significantly increased levels of percent tail DNA, indicative of an increase in damaged DNA (Fig. 2B). These results also correlate well with the IF data for *γ*H2AX, RPA32, and 53BP1 that show lower MFI for Vpr-induced DNA damage markers when compared to etoposide treatment (Fig. 1B). We segregated the samples into two populations, below 20% and above 20% tail DNA, to highlight the population of cells within each sample with a greater extent of damage (Fig. 2A and 2C). Whereas approximately 1% of uninfected and empty vector control cells had tail DNA above 20%, HIV-1 and HIV-2 Vpr expression resulted in 5% and 8% of cells above 20% tail DNA, respectively, indicating that expression of Vpr leads to significant DNA damage.

**Figure 2.**
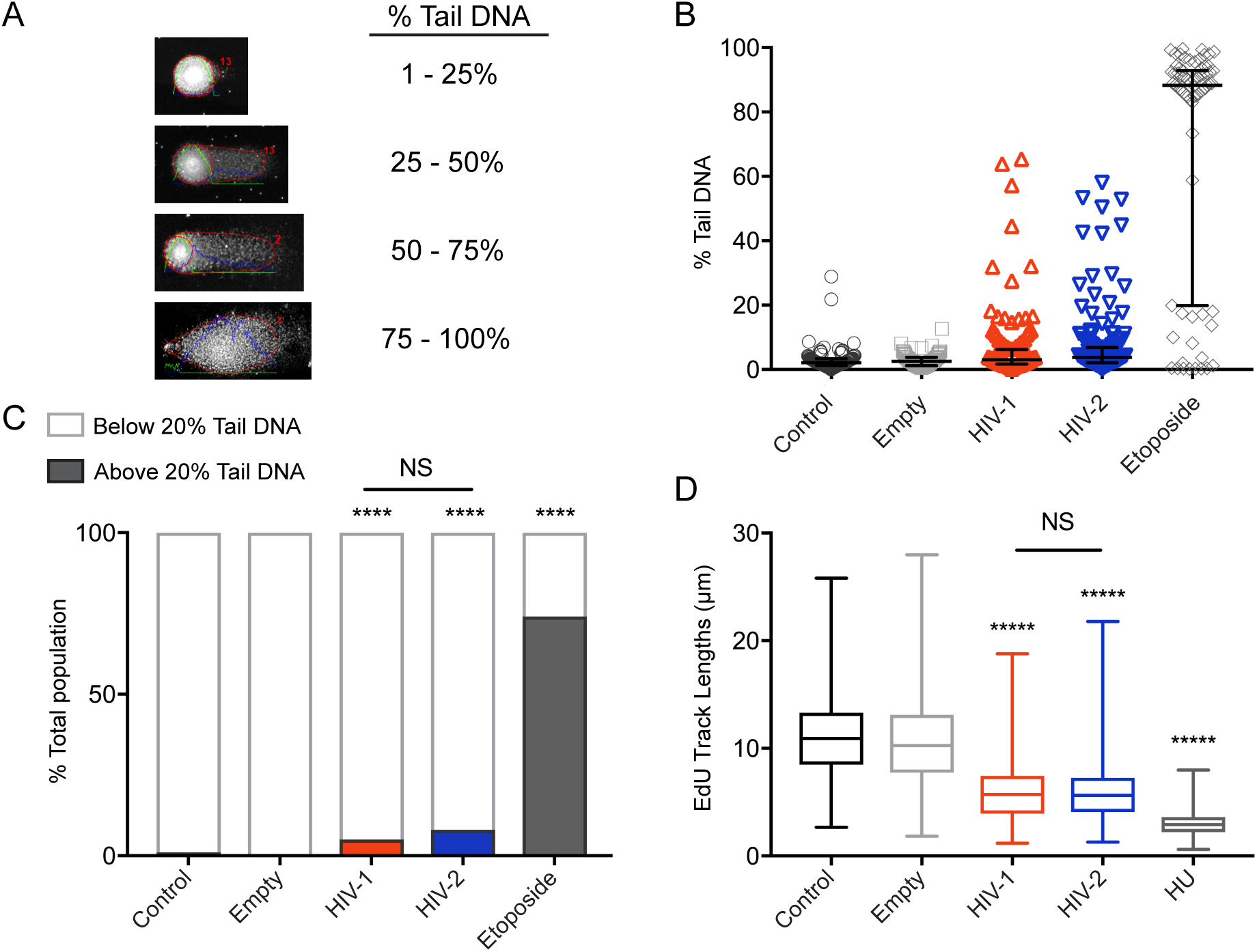
HIV-1 and HIV-2 Vpr damage DNA and stall DNA replication. A. Visual representation taken from the alkaline comet assay of the four varying degrees of damage measured by percent (%) tail DNA. Intensity profiles, lines, and numbers on the images were automatically generated by the *OpenComet* plug-in for the ImageJ software. B. Distribution of the % tail DNA measured for 100 cells per condition from one independent experiment using the *OpenComet* plug-in. U2OS cells were treated in the same conditions as Fig. 1A prior to being harvested for the comet assay. The bars represent the median with interquartile range; n=3, one representative experiment shown. C. A bar graph representation of the cells in panel B separated into two populations, below 20% tail DNA (unshaded) and above 20% tail DNA (shaded). Asterisks indicate statistical significance in comparison to empty vector control, or HIV-1 compared to HIV-2 Vpr, as determined by Chi Square test (NS, nonsignificant; *, *P*<0.05; **, *P*<0.01; ***, P<0.001; ****, *P*<0.0001; n=3, one representative experiment shown). D. A box and whiskers representation of the distribution of EdU track lengths (µm). U2OS cells were treated in the same conditions shown in Fig. 1A. Asterisks indicate statistical significance for HIV-1, HIV-2, and hydroxyurea (HU) compared to the empty vector control as determined by the Kruskall-Wallis test, while statistical difference between HIV-1 and HIV-2 was determined by the Mann-Whitney test (NS, nonsignificant; *, *P*<0.05; **, *P*<0.01; ***, P<0.001; ****, *P*<0.0001; n=3, one representative experiment shown).

As replication stress has been proposed to be a driver of this Vpr-induced DDR (51), and the activation of the DNA damage markers and cell cycle arrest (Fig. 1A and Fig. S1) are hallmarks of stalled DNA replication forks, we next determined whether Vpr expression leads to replication fork stalling via the DNA combing assay (52). This assay quantities the length of replication tracks by incorporation of EdU into nascent DNA. U2OS cells were infected with rAAV-Vpr for 20 hours, at which point EdU was added to the cells for 20 minutes. Hydroxyurea (HU), which stalls DNA replication by depleting dNTP pools (53), was used as a positive control. We found that HIV-1 and HIV-2 Vpr significantly decreased EdU track lengths compared to the uninfected and empty vector controls (Fig. 2D). Consistent with DNA damage markers, there was no direct correlation between levels of Vpr expression and DNA replication during this 20-minute window. However, cells expressing the highest levels of Vpr were largely not in S-phase during this window (Fig. S3), suggesting there is a threshold where Vpr expression robustly excludes cells from S-phase. Like the comet and IF assays, the greatest amount of replication fork stalling was exhibited by the positive control, HU, suggesting that while the impairment of normal DNA replication by HIV-1 and HIV-2 Vpr is significant, it is not as detrimental to the cell as HU. Overall, our alkaline comet and DNA combing data show that Vpr directly engages the DDR by inducing DNA breaks and stalling DNA replication.

### ATR senses stalled replication forks downstream of Vpr-induced DNA damage

Our results indicate that Vpr directly damages DNA and stalls DNA replication (Fig. 2). However, whether DNA damage occurs prior to replication fork stalling or as a consequence of stalled replication forks is unclear. To differentiate between these two possibilities, we inhibited the fundamental DNA damage repair kinase ATR via the selective ATR inhibitor (ATRi) VE-821 (54). ATR acts as the primary signaling axis for replication stress and cell cycle checkpoints, where it is recruited during S phase through RPA to stalled replication forks (9, 54). Here, it stabilizes replication forks from collapse, initiates the recruitment of repair proteins, and activates critical cell cycle checkpoints (9, 54). If Vpr-mediated DNA damage is due to stalled replication, we would expect ATR inhibition to increase DNA damage as the cells would not be able to guard against replication fork collapse or initiate repair. However, if damage occurs before replication stress, we would expect the inhibition of ATR to alter fork progression, but not DNA damage.

We first confirmed ATR inhibition mitigated Vpr-mediated cell cycle arrest for both HIV-1 and HIV-2 Vpr isolates tested (Fig. S4A). We also assayed for an effect of ATM inhibition (ATMi – KU-55933), as we found activation of repair markers associated with ATM activation (such as γH2AX and 53BP1 in Fig. 1), but found no effect of ATMi on Vpr-mediated cell cycle arrest (Fig. S4B), consistent with previously published results (32, 49, 55). Next, to determine the effect of ATR inhibition on DNA damage by Vpr, we again used the alkaline comet assay. While all samples had proportionately increased levels of damage when ATR was inhibited, there was no significant difference for either HIV-1 or HIV-2 Vpr with or without ATRi (Fig. 3A and 3B). This suggests that ATR inhibition does not affect the ability of Vpr to generate DNA lesions.

**Figure 3.**
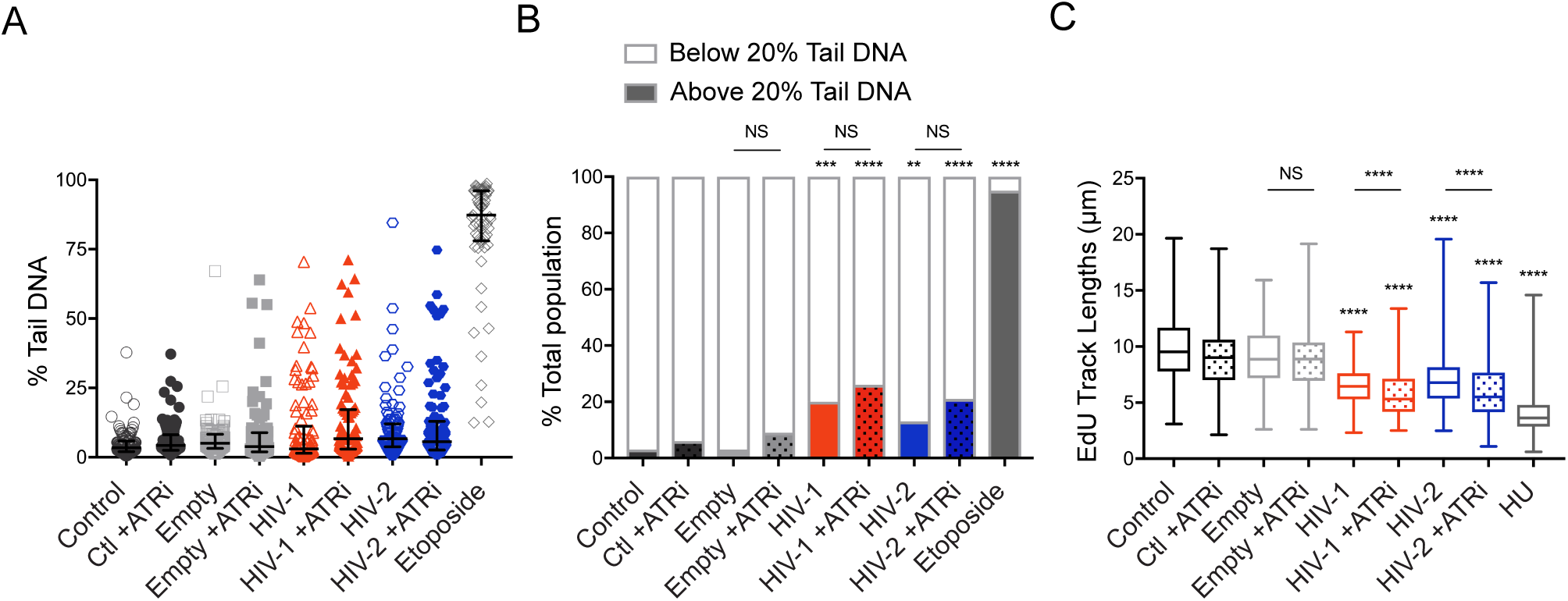
Vpr-induced DNA damage occurs prior to replication fork stalling and is independent of ATR. A. U2OS cells treated in the same conditions as Fig. 1A were incubated with or without 10µM VE-821 ATR inhibitor (ATRi) for 20 hours, then subjected to the alkaline comet assay as described in Fig. 2B. Graph shows quantification of % tail DNA of 100 cells measured per condition, with the bars representing the median and interquartile range. ATRi treated conditions shown in filled shapes (n=3, one representative experiment shown). B. A bar graph representation of the data from Fig. 3A, with the population separated as shown in Fig. 2C. Cells treated with ATRi above 20% tail DNA are represented as the shaded regions with dots. Asterisks indicate statistical significance as determined by Chi-square test (NS, nonsignificant; *, *P*<0.05; **, *P*<0.01; ***, P<0.001; ****, *P*<0.0001; n=3, one representative experiment shown). C. Distribution of EdU track lengths (µm) from cells treated with the same conditions as panel A. Cells treated with ATRi are represented as box plots with dots. Asterisks indicate statistical significance of empty vector +ATRi, HIV-1 +/− ATRi, HIV-2 +/− ATRi, and etoposide compared to empty vector -ATRi as determined by the Kruskall-Wallis test, while statistical difference between empty vector, HIV-1, and HIV-2 +/− ATRi is determined by the Mann-Whitney test (NS, nonsignificant; *, *P*<0.05; **, *P*<0.01; ***, P<0.001; ****, *P*<0.0001; n=3, one representative experiment shown).

In contrast, the DNA combing assay, which we used to determine the effect of ATR inhibition on stalled replication fork progression by Vpr, showed that replication track lengths were significantly shorter for HIV-1 and HIV-2 Vpr expressing cells when ATR was inhibited (Fig. 3C), presumably due to fork collapse. Though the overall effects of ATRi are modest, which is likely due to the intertwined nature of DNA damage, sensing, and repair, our data from the comet and DNA combing assays show that while Vpr mediated DNA damage is independent of ATR signaling, the ability to stall DNA replication is not. Moreover, it indicates that Vpr first induces DNA damage, which leads to the activation of ATR and subsequent stalled replication forks, presumably to mitigate replication stress.

### Vpr sensitizes cells to additional double-strand breaks

As we established with the immunofluorescence and alkaline comet assay, HIV-1 and HIV-2 Vpr induce DNA damage activating markers related to a wide variety of DNA lesions, such as SSB and DSB. While our data suggest that Vpr directly damages DNA, it is also possible that damage results from the inability of cells to repair pre-existing damage, such as damage due to replication stress. To address this question, we tested the sensitivity of cells expressing Vpr against various chemotherapeutics that directly damage DNA or inhibit a repair mechanism to cause damage.

We began by testing the sensitivity of Vpr treated cells to etoposide, which generates DSB by preventing the enzyme topoisomerase II from properly removing knots formed from DNA over-winding (56). Cells expressing Vpr were highly sensitized to etoposide treatment, where survival at even the lowest concentration (0.01uM) decreased to 60-70% compared to uninfected and empty vector control cells (Fig. 4). This indicates that Vpr expressing cells are unable to repair etoposide-induced DSB. We next tested sensitivity to hydroxyurea (HU). Prolonged exposure of HU to cells at high concentrations results in replication fork collapse and extensive DSB (57). Although Vpr expressing cells were not sensitized to HU treatment at low concentrations, at higher concentrations of HU (>3.90uM) where DSB are presumably present, survival of cells expressing HIV-1 and HIV-2 was significantly decrease compared to control cells (Fig. 4). Similar results were seen for the PARP1/2 inhibitor, Olaparib, which also leads to DSB due to the inability to repair DNA lesions (58) (Fig. 4). In contrast to the other chemotherapeutics, HIV-1 and HIV-2 Vpr expression did not dramatically hyper-sensitize cells to the interstrand-crosslinking agent cisplatin (59) (Fig. 4) despite activating markers associated with interstrand-crosslink (ICL) repair (6, 34). Altogether, the sensitivity assays indicate that Vpr expressing cells specifically show increased sensitivity to multiple chemotherapeutics that are capable of generating DSB by inhibiting crucial host repair mechanisms, suggesting that Vpr may also inhibit the ability of cells to repair this damage.

**Figure 4.**
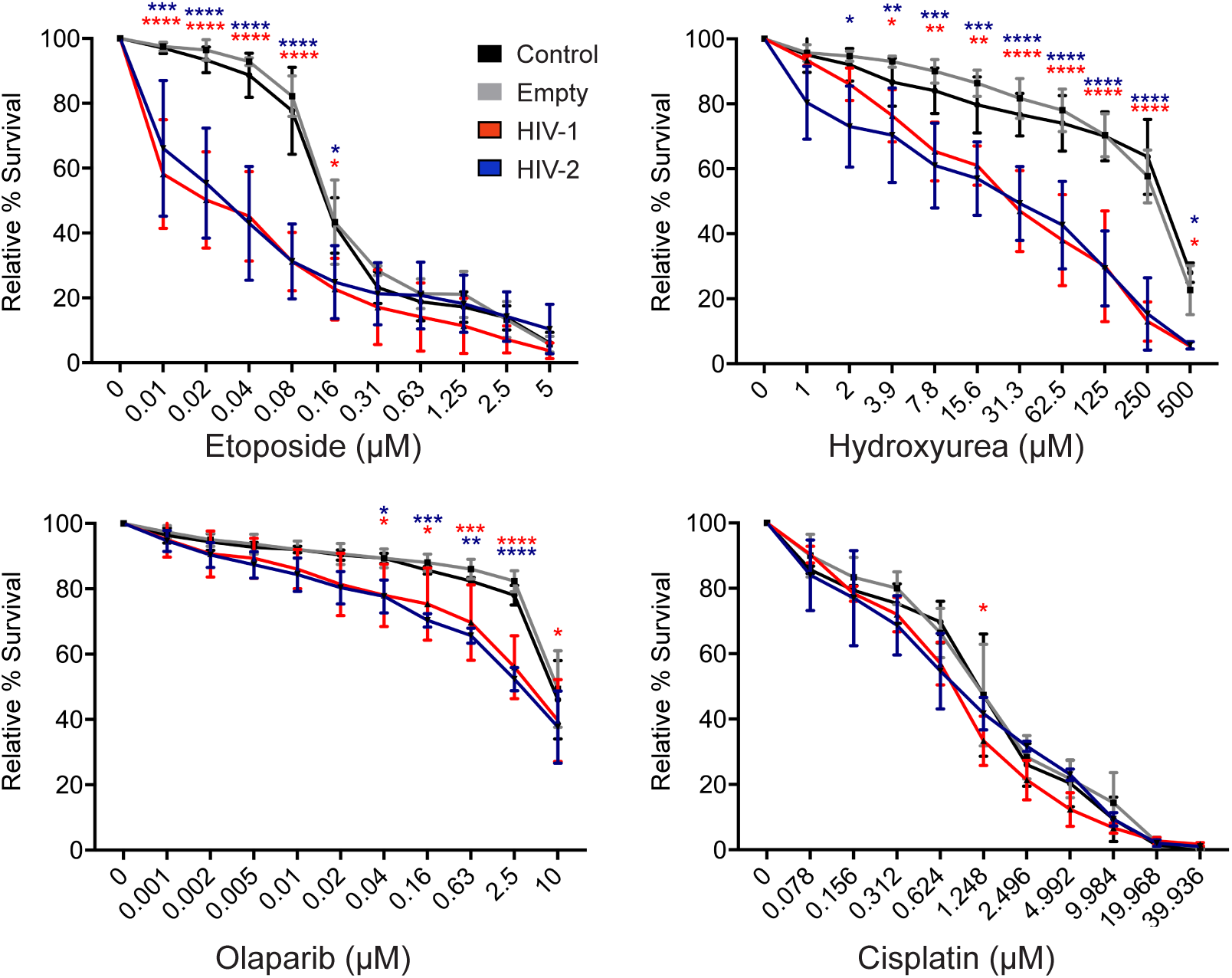
Cells expressing HIV-1 or HIV-2 Vpr are hypersensitive to exogenous double-strand DNA breaks. Sensitivities of the untreated control, empty vector control, HIV-1, and HIV-2 Vpr expressing U2OS cells to etoposide, hydroxyurea, olaparib, and cisplatin were tested by incubating cells for 7 days in the corresponding drug at the indicated concentrations. Survival was analyzed by crystal violet staining for live cells compared to the no drug treatment. Sensitivity results are the means of three independent experiments (n=3), error bars represent +/− standard deviations. Asterisks indicate statistical significance in comparison to empty vector control as determined by 2way ANOVA (*, p<0.05; **, p<0.03;***, p<0.002; ****, p<0.0001).

### Vpr inhibits double-strand break repair

Because we observed that Vpr expressing cells display hypersensitivity to the induction of exogenous DSB, we hypothesized that Vpr itself inhibits DNA break repair. To test this hypothesis, we used multiple independent GFP-based U2OS reporter cell lines that specifically monitor repair of an I-SceI induced DSB by either homologous recombination (HR), non-homologous end joining (NHEJ), alternative NHEJ (alt-NHEJ), or single-strand annealing (SSA) (60, 61). Each cell line contains a GFP gene that is uniquely disrupted by an I-SceI restriction site and does not express GFP, as well as a truncated GFP donor sequence. Upon transfection and expression of I-SceI, this site is cut and only proper repair by the indicated pathway results in GFP expression (see Fig. 5A and 5B for schematic of HR and NHEJ cell lines, respectively). In addition to transfecting I-SceI alone, we also used combinations that included empty vector, HIV-1, or HIV-2 Vpr that express mCherry via a T2A ribosomal skipping sequence. Thirty hours later, we measured repair on a per-cell basis using flow cytometry for successful repair (GFP) and transfection efficiency (mCherry) (Fig. 5A and 5B).

**Figure 5.**
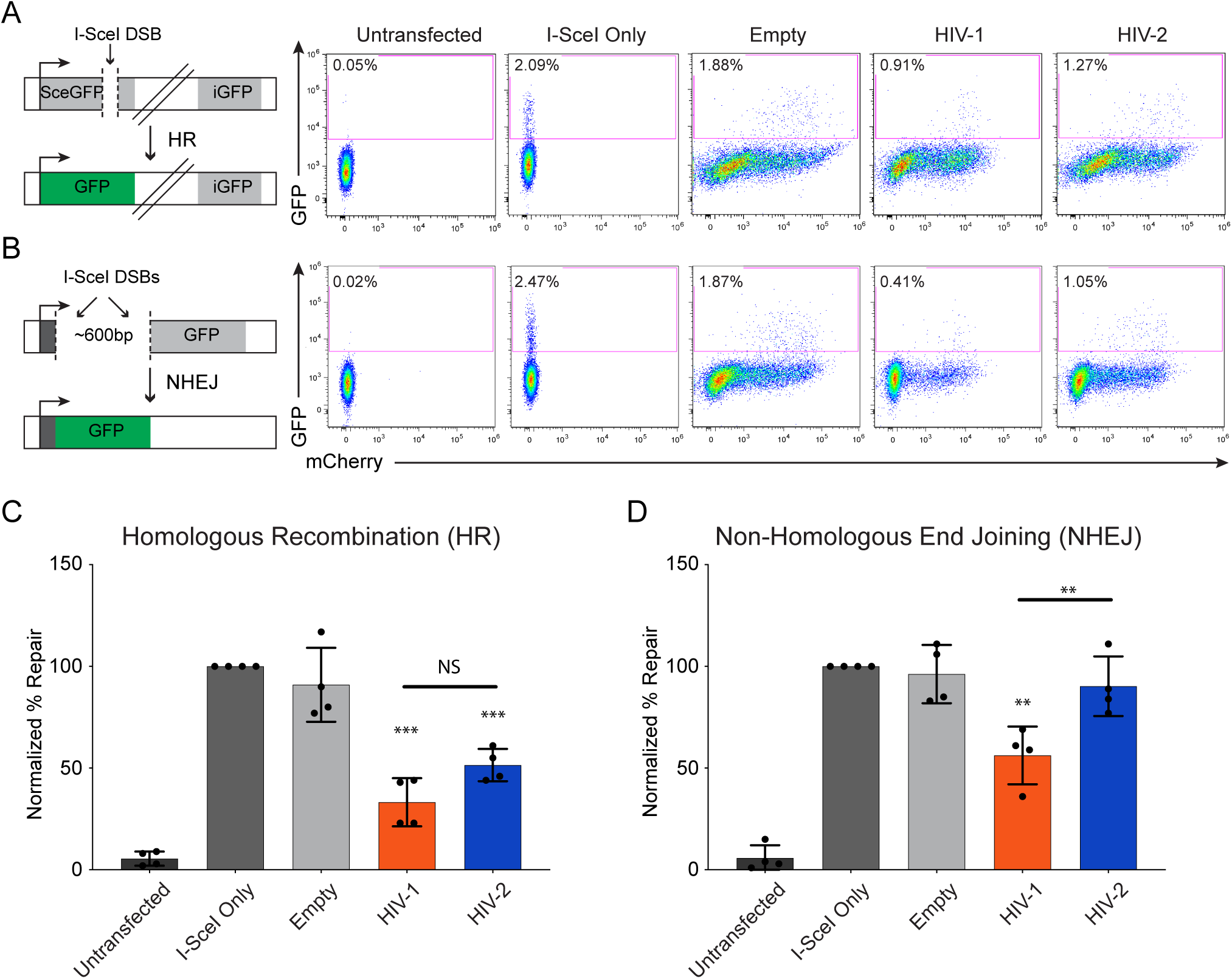
HIV-1 and HIV-2 Vpr repress double-strand break repair. A. (Left) Schematic of I-SceI-based homologous recombination (HR) U2OS reporter cell line (“DR-GFP” assay). (Right) Representative flow cytometry plots of one I-SceI repair assay experiment for HR repair. Cells were transfected for 30 hrs with the I-SceI plasmid alone or with either empty vector, HIV-1, or HIV-2 Vpr that expresses mCherry via a T2A ribosomal skipping sequence. 20,000 cells were measured per condition and gated for homologous recombination-mediated DSB repair (GFP) B. (Left) Schematic of I-SceI-based classical non-homologous end joining (NHEJ) U2OS reporter cell line (“EJ5-GFP” assay). (Right) Representative flow cytometry plots of one I-SceI repair assay experiment for NHEJ repair. Cells were treated and measured in the same conditions as described in panel A. C. I-SceI HR repair assay representing the average percent repair by homologous recombination from four experiments (n=4), normalized to the I-SceI Only condition. Cells were treated and measured using the same conditions described in panel A. Asterisks indicate statistical significance compared to empty vector control as determined by a one sample t test (theoretical mean set to the average value of the empty vector control), while statistical difference between HIV-1 and HIV-2 was determined by the Mann-Whitney test (NS, nonsignificant; *, *P*<0.05; **, *P*<0.01; ***, P<0.001; ****, *P*<0.0001). Error bars represent +/− standard deviation D. I-SceI NHEJ repair assay representing average percent repair by classical non-homologous end joining from four experiments (n=4), normalized to the I-SceI Only condition. Cells were treated and measured in the same conditions as described in panel A. Statistical analysis was determined with the same methods as shown in panel B. Error bars represent +/− standard deviation.

We first tested the I-SceI reporter cell line for HR. While transfection of I-SceI alone or with empty vector control resulted in similar amounts of HR, we found that cells transfected with HIV-1 and HIV-2 Vpr decreased HR efficiency by 66% and 49%, respectively, when normalized to control cells at 100% (Fig. 5A and 5C). This indicates that HIV-1 and HIV-2 Vpr repress HR. Based on these results, we next tested the I-SceI reporter cell line that measures NHEJ, which is often utilized by cells to repair DSBs when HR is repressed (62). Similar to HR, HIV-1 Vpr expression also decreased NHEJ efficiency by 51% compared to wild type cells. In contrast to HIV-1, HIV-2 Vpr did not significantly decrease NHEJ, as these cells were able to repair via NHEJ at 90% of wild type levels (Fig. 5B and 5D), highlighting potential mechanistic differences between HIV-1 and HIV-2 Vpr. And consistent with DNA damage and DNA replication, there was no correlation between Vpr expression (mCherry) and repair (GFP) based on flow plots (Fig. 5A and 5B). Finally, we tested the I-SceI reporter cell lines for alt-NHEJ and SSA repair mechanisms, but found no significant change in repair when compared to control cells (Fig. S5). Thus, based on the data from the four different I-SceI reporter cell lines, we have identified that both HIV-1 and HIV-2 Vpr repress double-strand break repair, in addition to inducing DNA damage.

### Disconnect between induction of DNA damage and downregulation of repair machinery

Our findings demonstrate that both HIV-1 and HIV-2 Vpr are capable of inducing DNA damage, stalling DNA replication, downregulating double-strand DNA break repair, and causing cell cycle arrest. However, it is unclear how these phenotypes are linked and what role(s) host protein interactions play. To address these questions, we further tested a subset of well characterized HIV-1 and HIV-2 Vpr mutants for their ability to induce, signal, and respond to DNA damage via the alkaline comet assay, EdU immunofluorescence, HR I-SceI repair assay, and bivariate cell cycle analysis, respectively. We tested four mutants for each HIV-1 and HIV-2 Vpr (Fig S6A). These include: HIV-1 W54R/HIV-2 L59A Vpr mutants, which block the ability of HIV-1 Vpr to recruit and degrade the DNA glycosylase UNG2 (63); HIV-1 Q65R/HIV-2 Q70R Vpr, which renders Vpr unable to properly localize, multimerize, or recruit known host proteins such as the Cul4A^DCAF1^ complex or UNG2 and is therefore largely functionally dead (33, 64, 65); HIV-1 S79A/HIV-2 S84A mutants, which renders Vpr unable to cause cell cycle arrest or interact with TAK1 to activate canonical NF-kB (66, 67); HIV-1 R80A/HIV-2 R85A Vpr mutants, which can still interact with Cul4A^DCAF1^ and degrade TET2 (41) but do not cause cell cycle arrest, presumably due to the requirement of an additional unknown host protein(s) (68). Moreover, as HIV-1 but not HIV-2 Vpr interacts with UNG2, HLTF, and the SLX4 complex (6, 37), by testing diverse Vpr orthologs we were further able to dissect the requirement(s) for previously reported Vpr-interacting proteins in inducing DNA damage, stalling DNA replication, downregulating HR repair, and causing cell cycle arrest.

Consistent with previously published results, all mutants except HIV-1 W54R/HIV-2 L59A Vpr failed to induce cell cycle arrest (Fig. 6A and Fig. S6B). In contrast to cell cycle arrest, only HIV-1 Q65R/HIV-2 Q70R Vpr lost the ability to damage DNA (Fig. 6B and Fig. S6C), indicating that damage of DNA occurs independent of cell cycle arrest and independent of the Vpr-host protein-protein interactions assayed here. When testing for the effects of Vpr on DNA replication, we found that, in addition to HIV-1 Q65R/HIV-2 Q70R Vpr, HIV-1 S79A/HIV-2 S84A Vpr mutants were unable to stall DNA replication (Fig. 6C), suggesting that activation of TAK1 is integral in the ability of Vpr to stall DNA replication. Finally, in concert with cell cycle arrest, all mutants except the HIV-1 W54R/HIV-2 L59A Vpr mutants failed to repress homologous recombination repair (Fig. 6D). A summary of these results can be found in Table 1. Overall, our mutational analyses of HIV-1 and HIV-2 Vpr indicate that repression of HR and cell cycle arrest are correlated, and that these two phenotypes are independent of Vpr-induced DNA damage and downstream signaling. Moreover, by testing multiple mutants deficient for host factor recruitment, as well as comparing HIV-1 and HIV-2 Vpr orthologs which differentially recruit host proteins, our results rule out most previously observed Vpr-interacting host proteins for a role in induction of DNA damage and repression of HR.

**Table 1.** Summary of mutant HIV-1 and HIV-2 Vpr data. Summary of data from Figure 6. Plus sign (+) indicates that Vpr was functional in the indicated assay, while minus sign (-) indicates that Vpr was statistically indistinguishable from empty vector control. The solid vertical bold line indicates potential separation of Vpr function based on mutant Vpr analysis.

|  | DNA<br>Damage | Replication<br>Stalling | HR<br>Repression | Cell Cycle<br>Arrest |
| --- | --- | --- | --- | --- |
| **HIV-1/HIV-2** | + | + | + | + |
| **W54R/L59A** | + | + | + | + |
| **Q65R/Q70R** | - | - | - | - |
| **S79A/S84A** | + | - | - | - |
| **R80A/R85A** | + | + | - | - |

**Figure 6.**
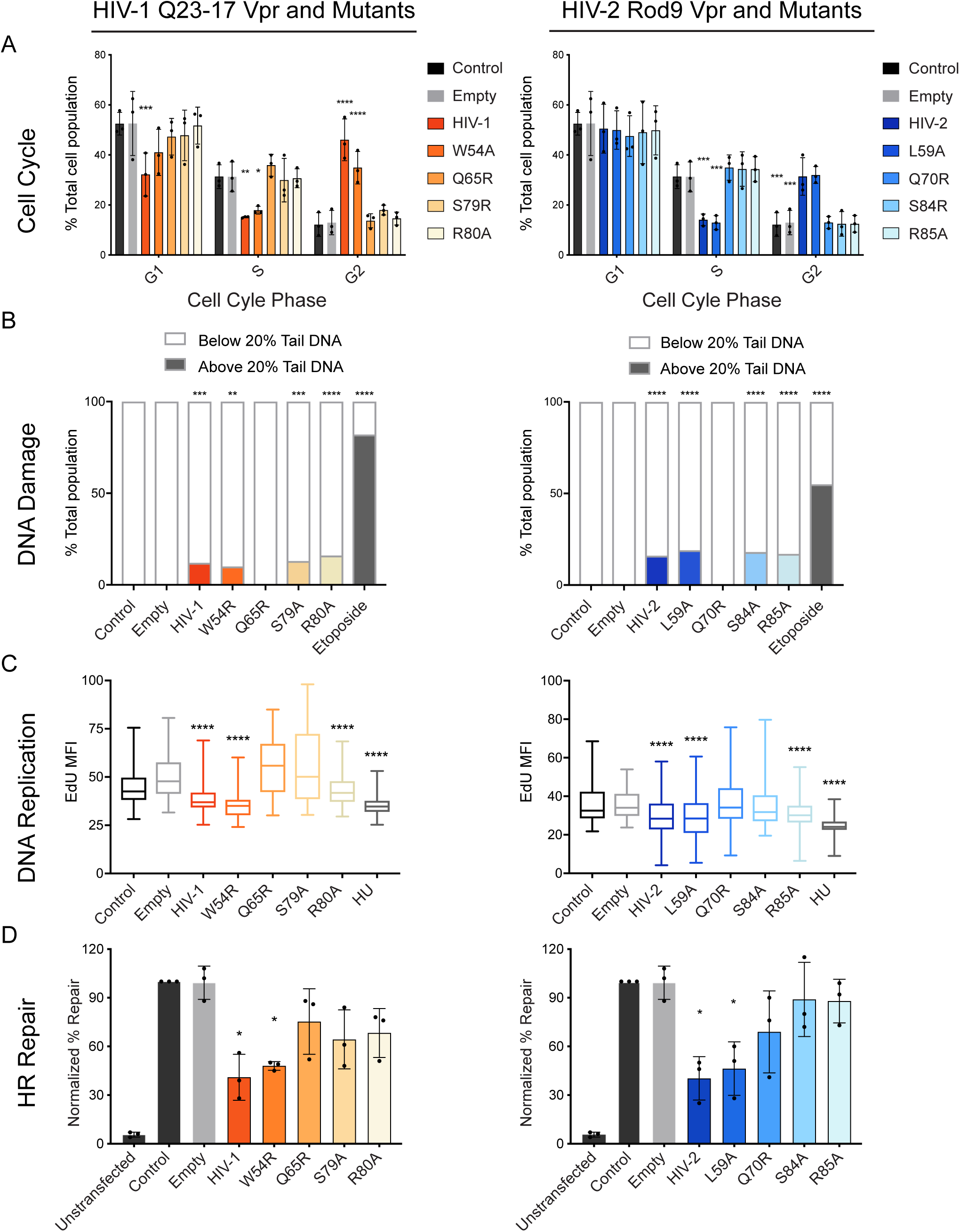
Vpr-induced DNA damage is independent of repression of homologous recombination and cell cycle arrest. A. Bivariate cell cycle analysis of synchronized U2OS cells infected with rAAV expressing 3X FLAG-tagged HIV-1 and HIV-2 Vpr, empty vector, or control uninfected cells for 38 hours. The graph shows the percentage of the population of 10,000 cells per condition in G1, S, and G2 measured using flow cytometry of cells stained for propidium iodide (PI, total DNA content) and EdU (DNA synthesis). Asterisks indicate statistical significance in comparison to empty vector control as determined by Tukey multiple comparison test (NS, nonsignificant; *, *P*<0.05; **, *P*<0.01; ***, P<0.001; ****, *P*<0.0001; n=3). Error bars represent +/− standard deviation. B. The alkaline comet assay for HIV-1 and HIV-2 Vpr mutants as represented in Fig. 2C with 100 cells measured per condition. U2OS cells were treated in similar conditions as Fig. 1A. Asterisks indicate statistical significance to empty vector control as described in Fig. 2C (n=3, one representative image shown). C. Box and whisker plot representation of the distribution of EdU mean fluorescence intensity (MFI) for HIV-1 and HIV-2 Vpr mutants with cells treated in the same condition as panel B. Asterisks indicate statistical significance to empty vector control as determined by the Dunn’s multiple comparison test (NS, nonsignificant; *, *P*<0.05; **, *P*<0.01; ***, P<0.001; ****, *P*<0.0001; n=3, one representative experiment shown). D. Experimental results from the I-SceI DR-GFP assay representing average percent repair by homologous recombination for HIV-1 and HIV-2 mutants as described in Fig. 5C. Asterisks indicate statistical significance from empty vector control as described in Fig. 5C (n=3).

### Repression of HR is not a consequence of Vpr-mediated cell cycle arrest

The predominant phenotype of Vpr expression *in vivo* and *in vitro* is G2 cell cycle arrest. While it is unclear what leads to Vpr-mediated cell cycle arrest, G2 arrest depends on recruitment of the Cul4A^DCAF1^ ubiquitin ligase complex through a direct interaction of Vpr with DCAF1. Here we have identified a new phenotype of Vpr, repression of HR, that tracks with G2 cell cycle arrest based on our Vpr mutant data (Fig 6 and Table 1). However, whether repression of HR by Vpr is a consequence or potential driver of Vpr-mediated arrest remains unclear.

To address this, we first asked if Cul4A^DCAF1^ complex recruitment is also required for repression of HR by Vpr. We selected two mutants that have been previously shown to alter HIV-1 Vpr binding to DCAF1, L64A (28) and H71R (35), and further generated those mutants in HIV-2 Vpr (L69A and H76R, respectively). To validate if these mutants lost the ability to recruit DCAF1, we immunoprecipitated FLAG-Vpr and probed for endogenous human DCAF1. In our hands, HIV-1 H71R/HIV-2 H76R no longer recruited DCAF1. However, HIV-1 L64A/HIV-2 L69A was still able to recruit the DCAF1 adaptor protein, though at a slightly lower level than wild type Vpr (Fig. 7A). Consistent with recruitment of DCAF1, HIV-1 H71R/HIV-2 H76R, but not HIV-1 L64A/HIV-2 L69A, fully lost their ability arrest cells (Fig. S7). We next tested these mutants for their ability to repress HR using the HR I-SceI repair assay. Again, consistent with DCAF1 binding and cell cycle arrest, HIV-1 H71R/HIV-2 H76R failed to repress HR, whereas HIV-1 L64A/HIV-2 L69A repressed HR to near WT Vpr levels (Fig. 7B). These data suggest that, similar to cell cycle arrest, repression of HR repair by Vpr requires DCAF1 binding.

**Figure 7.**
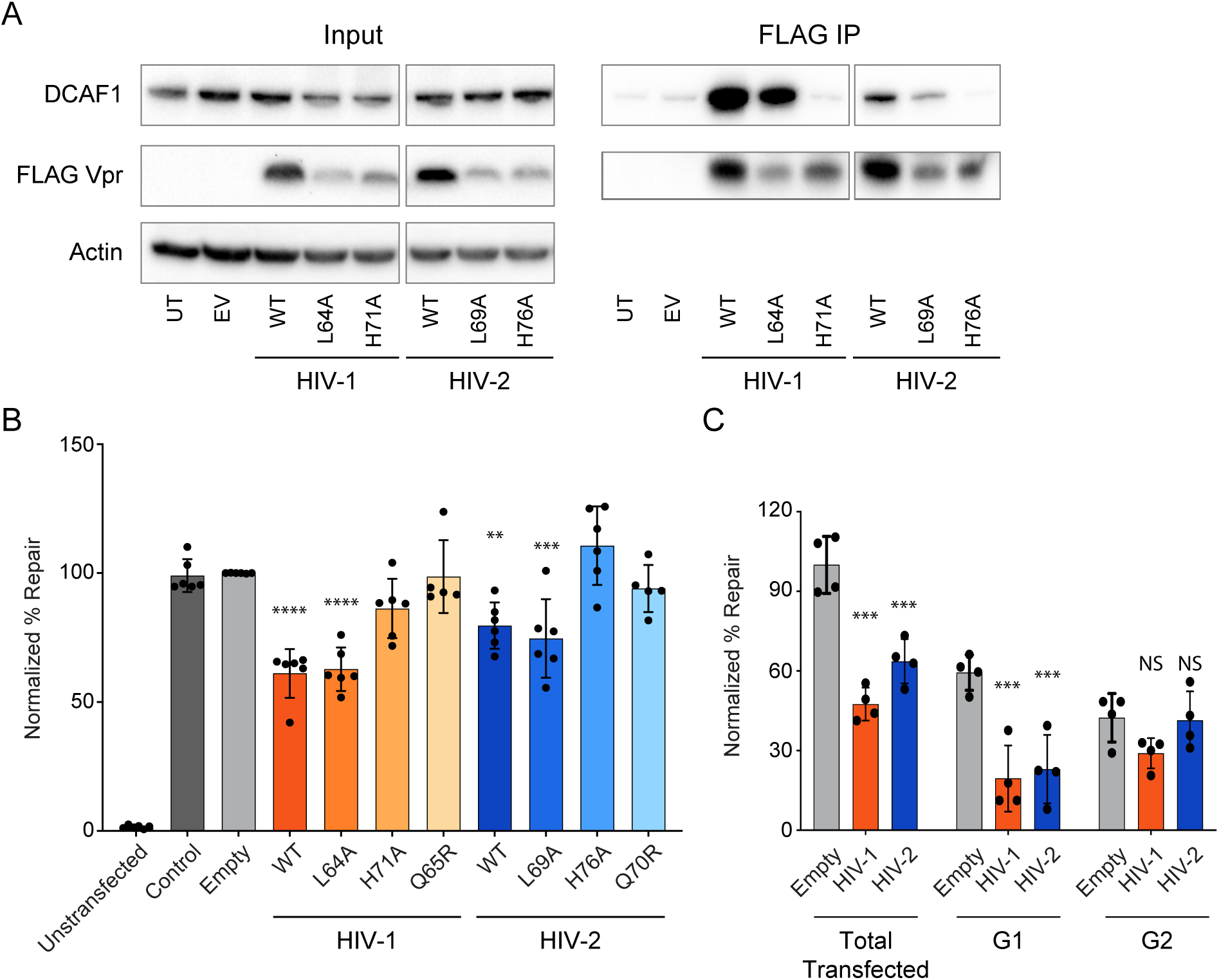
Repression of homologous recombination by Vpr requires DCAF1 and precedes cell cycle arrest. A. Representative western blots of U2OS cells for endogenous DCAF1, transiently transfected 3X-FLAG Vpr, and endogenous actin as a loading control (left “Input” panels). Immunoprecipitations against 3X-FLAG, probed for endogenous DCAF1 and transiently transfected 3X-FLAG Vpr (right “FLAG-IP” panels). B. Experimental results from the I-SceI DR-GFP assay representing average percent repair by homologous recombination for HIV-1 and HIV-2 mutants as described in Fig. 5C. Asterisks indicate statistical significance from empty vector control as described in Fig. 5C (n=6). C. Experimental results from bivariate I-SceI DR-GFP-cell cycle assay. Cells were transfected for 30 hrs with the I-SceI plasmid alone or with either empty vector, HIV-1, or HIV-2 Vpr that expresses mCherry via a T2A ribosomal skipping sequence, then labeled with Hoechst dye to label total DNA content. 20,000 cells were measured per condition. Total, G1, and G2 mCherry expressing cell populations were gated for homologous recombination-mediated DSB repair (GFP). Asterisks indicate statistical significance from empty vector control as described in Fig. 5C (n=4).

To determine if repression of HR by Vpr requires G2 arrest or occurs independent of this arrest, we defined the cell cycle status (G1 or G2 phase) of DR-GFP cells that exhibited repair using Hoechst dye. We would expect that if Vpr-mediated G2 arrest is required to repress HR, then Vpr-expressing cells in G2 would primarily show repressed HR. However, if G2 arrest is not required for Vpr to repress HR, then cells in G1 would also show a repression of HR in the presence of Vpr.

As seen previously, both HIV-1 and HIV-2 Vpr repressed total cellular HR when compared to empty vector control. Vpr-expressing cells also showed strong repression of HR repair in G1 when compared to empty vector control cells. However, Vpr-expressing cells did not repress HR in G2, as they were statistically indistinguishable from control cells (Fig. 7C). Together, this data indicates that Vpr-mediated repression of HR does not require G2 arrest, but instead occurs primarily in the G1 phase of the cell cycle. Moreover, as G1 precedes G2, this may suggest that repression of HR is the initiating step leading to cell cycle arrest, and therefore may be the crucial phenotype associated with the primary function of Vpr.

## DISCUSSION

Here we show that HIV-1 and HIV-2 Vpr induce both double- and single-strand DNA breaks, leading to the recruitment of repair factors, including *γ*H2AX, RPA32, and 53BP1. These Vpr-induced DNA lesions are sensed by ATR and require NF-kB signaling to stall DNA replication. However, contrary to the induction of DNA damage and the activation of the DNA damage response, Vpr represses essential mechanisms of double-strand break repair, including homologous recombination repair (HR) and non-homologous end joining (NHEJ). Mutational analysis of Vpr has identified that there is a disconnect between mutants that can damage DNA and those that can repress DNA repair and activate cell cycle arrest. Finally, we show that repression of HR is not a consequence of G2 cell cycle arrest, but instead is potentially a driver of this arrest. Overall, our data support a model where Vpr has two unique and independent mechanisms to modulate the host DDR: first, Vpr has the inherent ability to induce DNA damage, which is largely independent of known Vpr-binding host factors. This Vpr-induced damage is sensed by ATR, and signals through NF-kB to block DNA replication fork progression. Second, through recruitment of the Cul4A^DCAF1^ complex, Vpr represses DNA double-strand break repair machinery, leading to a prolonged cell cycle to deal with the inability to repair DNA lesions.

Why would Vpr engage the DDR at two unique steps, and how would this help lentiviral replication? While it may seem counterintuitive to both activate and repress the DDR through unique mechanisms, Vpr is not the only viral protein, and lentiviruses are not the only viruses, to both activate and repress the DDR at different steps in viral replication (14). For example, human papillomaviruses (HPV) upregulate ATM in order to push cells away from NHEJ and towards HR, which is thought to enhance viral persistence and integration (69, 70). Interestingly, this also sensitizes HPV+ cells to exogenous genotoxic agents due to their inability to repair additional damage (71), as we have shown here for HIV Vpr (Figure 4). Moreover, as Vpr has two unique phases in an infected cell – it is delivered early via the incoming virion and expressed *de novo* following integration and gene expression – it is possible that these two distinct DDR-associated functions of Vpr are separated in the viral lifecycle of an infected cell.

While it is possible that some of these DDR-associated phenotypes are indirect consequences of other effects of Vpr on the cell, such as induction of pro-inflammatory cytokines (72), this dual function of Vpr in engaging the DDR at multiple independent steps could potentially help clarify some of the discrepancies in the Vpr literature, and may directly explain many of the roles in viral replication attributed to Vpr (73–79). For example, DNA damage promotes nucleotide biosynthesis (80) and thus may enhance early events in HIV replication such as reverse transcription. This is analogous to the degradation of SAMHD1 by lentiviral Vpx/Vpr (5, 81, 82) and could help to explain why Vpr from HIV-2, which encodes both Vpr and the paralogous Vpx protein, does not attenuate host repair machinery, or recruit host DDR proteins (6, 36, 37, 40, 41, 83), as efficiently as HIV-1 Vpr. The stalling of replication forks (Fig 2D) could enhance integration by remodeling histones and prolonging S-phase. Integration could also be enhanced by attenuating double-strand break repair (Fig. 5), similar to the repression of HR and base excision repair by Human T-lymphotropic Virus 1 (HTLV-1) to facilitate viral integration (84–86). Moreover, the induction of DNA breaks (Fig. 2A-C) could enhance LTR-driven transcription by activating important DDR-responsive transcription factors, such as NF-kB and AP-1 (67, 87).

As the primary role of lentiviral accessory genes is to overcome antiviral restriction factors, our data also support a model where DDR proteins and/or pathways restrict HIV replication and are overcome by Vpr. This is consistent with the growing evidence that DDR proteins and pathways contribute to the innate immune response to response to pathogens (17–22). We have shown that, like Vpr-mediated cell cycle arrest, recruitment of the Cul4A ubiquitin ligase complex adaptor protein DCAF1 is required for repression of HR repair (Fig. 7). Vpr could be recruiting this complex away from a natural target, or usurping it to degrade a host protein, which is consistent with the primary role of lentiviral accessory genes in viral replication, such as Vpx-mediated degradation of the antiviral DDR protein SAMHD1 (88, 89). And while Vpr has been shown to recruit and degrade many host proteins, through the combination of our mutant data and use of HIV-1 and HIV-2 Vpr orthologs (Fig. 6 and 7), we are able to rule out most known DDR-associated Vpr-interacting proteins (and potential cellular effects of Vpr) for roles in modulating the DDR as described herein. Whether some of the remaining Vpr-interacting proteins we were unable to characterize, such as the endonuclease Exo1, are required for Vpr-mediated engagement of the DDR, or whether novel undiscovered host proteins are required remains unclear. Moreover, whether modulation of DDR pathways is a direct primary effect of Vpr or a consequence of degradation of an antiviral host protein that is also integral to the DDR is also unclear. However, our data pinpoint double-strand DNA break repair as important cellular pathways that warrant further investigation into both innate immunity and Vpr.

Our mutant data also shows that the long-standing enigmatic cell cycle arrest caused by Vpr correlates with repression of HR, suggesting these two phenotypes are linked. As HR is upregulated in G2, one might expect Vpr to enhance this repair mechanism instead of inhibit it. Intriguingly, we find the majority of Vpr-mediated repression of HR occurs in cells that are currently in G1, not G2. This indicates that repression of HR precedes G2 arrest, and is the initiating step which ultimately leads to G2 arrest. Based on this, we hypothesize that repression of HR, not cell cycle arrest, is the crucial phenotype associated with Vpr, and that understanding this process will give clearer insight into the primary function of Vpr in viral replication.

Thus, while it is clear that the DDR is a central hub that is essential for replication of many viruses in different phases of their lifecycle, the precise roles of Vpr-mediated activation and repression of the DDR in HIV replication remain obscure. In establishing that Vpr activates and represses the DDR, we have clarified the multiple ways that Vpr modulates the host DDR and uncovered a new phenotype for Vpr that may precede cell cycle arrest, suppression of double-strand break repair. This will allow us to better define the primary evolutionarily-conserved role of Vpr. Finally, our data indicate that Vpr expression could have important implications for the development and treatment of HIV-associated diseases such as cancer, where induction of DNA damage and deregulation of repair could serve to complicate tumorigenesis but also sensitize cells to chemotherapeutics; further highlighting the importance of Vpr in HIV replication and associated diseases.

## METHODS

### Plasmids

pscAAV-mCherry-T2A-Vpr plasmids were generated by replacing GFP with mCherry from pscAAV-GFP-T2A-Vpr (6). HIV-2 A.PT (A.PT.x.ALI.AF082339) and HIV-2 G.CI.92 (G.CI.92.Abt96.AF208027) were synthesized as gBlocks (IDT) and subcloned into the pscAAV-mCherry-T2A-Vpr construct using standard cloning techniques. Vpr mutants were generated using site directed mutagenesis (Q5 site directed mutagenesis kit, NEB). pCBASceI was a gift from Maria Jasin (Addgene plasmid # 26477) (90).

### Cell lines & cell culture

Human embryonic kidney (HEK) 293, HEK 293T, and human bone osteosarcoma epithelial (U2OS) cells were cultured as adherent cells directly on tissue culture plastic (Greiner) in DMEM growth medium (high glucose, L-glutamine, no sodium pyruvate; Gibco) with 10% fetal bovine serum (Gibco) and 1% penicillin-streptomycin (Gibco) at 37°C and 5% CO_2_. All cells were harvested using 0.05% Trypsin-EDTA (Gibco). Transfections were performed with TransIT-LT1 (Mirus). The panel of U2OS cells containing an integrated reporter (DR-GFP, SA-GFP, EJ2-GFP, EJ5-GFP) used in the I-SceI repair assays were kindly provided by Jeremy M. Stark (Beckman Research Institute of the City of Hope) (60).

### Generation of Viruses

AAV vectors were generated by transient transfection of HEK 293 cells using polyethyleneimine (PEI) as previously described (91). Levels of DNase-resistant vector genomes were quantified by inverted terminal repeat (ITR)-specific quantitative PCR (qPCR) using a linearized plasmid standard according to the method of Aurnhammer et al. (92).

### Western Blots & co-immunoprecipitations

Cells we lysed in radioimmunoprecipitation assay (RIPA) buffer (50 mM Tris-HCl [pH 8.0], 150 mM NaCl, 1 mM EDTA, 0.1% SDS, 1% NP-40, 0.5% sodium deoxycholate, Benzonase, Protease inhibitor), and clarified by centrifugation at 14,500g for 10 min. Immunoprecipitations were performed as previously described (6) using anti-FLAG affinity beads (Sigma). All samples were boiled in 4X sample buffer (40% glycerol, 240mM Tris pH 6.8, 8% SDS, 0.5 % β-mercaptoethanol, and bromophenol blue) in preparation for SDS-PAGE using 4-12% Bis-Tris polyacrylamide gels and subsequently transferred onto a PVDF membrane. Immunoblotting was performed using: mouse anti-FLAG (Sigma), mouse anti-Actin (Thermo-Fisher), rabbit anti-DCAF1 (Cell Signaling) goat anti-mouse HRP (Invitrogen), goat anti-rabbit HRP (Invitrogen).

### DNA Combing Assay

DNA combing assay was adapted from (52). Cells were plated in 6-well tissue culture treated plates (Greiner) at 1.75 × 10^6^ cell/ well and allowed to rest overnight. Cells were then infected with rAAV 2.5 at equal titers (1.4 × 10^8^ copies/ well) or 500uM hydroxyurea (Sigma) for 20 hrs. Following infection, cells were incubated with 10uM EdU (Invitrogen) for 20 min., then harvested, spun down, and resuspended in 1X PBS (Gibco). The cell suspension was added and lysed with lysis buffer (50mM EDTA, 0.5% SDS, 200mM Tris-HCl pH 7.5) directly on a silane-coated slide (Electron Microscopy), then incubated for 5 to 8 min. After incubation, the slide was tilted at a 45° angle to allow the droplet to roll down then fixed with 3:1 methanol acetic acid for 15 min after the slide was completely dry. Then the slide was washed with 1X PBS, blocked with 3% BSA for 30 min, and stained with secondary EdU mixture (Click-IT EdU imaging kit; Invitrogen) and DNA (Yoyo-1; Life Technologies). Microscopy was performed using the Zeiss Axioimager Z1 and images were analyzed using ImageJ.

### Alkaline Comet Assay

Alkaline Comet Assay was performed as previously described (50) with some minor changes. Cells were plated in 6-well tissue culture treated plates (Greiner) at 1.75 × 10^6^ cell/ well and allowed to rest overnight. Cells were then infected with rAAV 2.5 at equal titers (1.4 × 10^8^ copies/ well) or 50uM etoposide (Sigma) for 20 hrs. Following infection, cells were then harvested, spun down, and resuspended in 0.5% low melting point agarose at 37°C. Samples were then spread onto agarose-coated slides (Cell Biolabs) and allowed to solidify for 20 min at 4°C. After agarose solidification, samples were incubated in lysis buffer (10 mM Tris-HCl pH 10, 2.5M NaCl, 0.1M EDTA, 1% Triton X-100) for 1 hr, then in the alkaline running buffer (0.3M NaOH, 1 mM EDTA) for 30 min., and finally electrophoresed at 300mA for 30min – all done at 4°C. Samples were then washed in ddH_2_O and fixed in 70% ethanol at 4°C. Cells were stained with Yoyo-1 (Life Technologies) for 15 min at room temperature, then washed with ddH_2_O and dried overnight. Images were acquired on the Zeiss Axioimager Z1. Images were analyzed using the OpenComet plug in for ImageJ.

### Cell Cycle Analysis

U2OS cells were plated and either left unsynchronized or synchronized using serum starvation with 0.05% FBS DMEM (Gibco) for at least 12 hours. Cells were infected with AAV2.5 (600 copies/cell) for 38 hours. For labeling with Hoechst, cells were incubated with Hoechst Ready Flow reagent (Invitrogen) as recommended. For labeling with propidium iodide, cells were fixed with ice-cold ethanol and DNA was stained with 0.01 g/ml propidium iodide (Sigma-Aldrich) and RNase A in PBS. For bivariate labeling, cells were additionally pulse labeled with 10 µM EdU (Invitrogen) for at least 30 minutes. Pulse labeled cells were then permeabilized with 0.01% Triton X-100 for 3.5 minutes and fixed with 4% PFA for 20 min. EdU was detected using Click-iT EdU Alexa Fluor 647 Imaging Kit (Invitrogen) followed by Hoechst or PI staining. Cells were assessed by flow cytometry on a FACSVERSE (BD). At least 10,000 cells were collected each run and data was analyzed using FlowJo software.

### Immunofluorescence

Cells were plated in 6-well tissue culture treated plates (Greiner) at 1.75 × 10^6^ cell/ well and allowed to rest overnight. Cells were then infected with rAAV 2.5 at equal titers (1.4 × 10^8^ copies/ well) or 50uM etoposide (Sigma) for 20 hrs. For the EdU-IF experiments, EdU was added to the cells for 20 minutes. Cells were then permeabilized with 0.5% Triton X-100 in PBS at 4°C for 5 min, fixed in 4% PFA for 20 min. Samples were then washed in 1X PBS and incubated with blocking buffer (3% BSA, 0.05% Tween-20, and 0.04 NaN3 in PBS) for 30 minutes. Cells were probed with appropriate primary antibodies (anti-FLAG M2 [Sigma-Aldrich], anti-γH2AX, anti-RPA32 [GeneTex], or anti-53BP1 [Cell Signaling]), then washed in PBST (0.05% Tween-20 in PBS), and probed with Alexa-Fluor conjugated secondary antibodies (Life Technologies). Nuclei were stained with diamidino-2-phenylindole (DAPI; Life Technologies). Secondary staining for EdU was added as the last step and stained twice to ensure signal. Images were acquired on the Zeiss Axioimager Z1 and mean fluorescence intensity (MFI) was analyzed using ImageJ.

### Sensitivity Assays

Sensitivity assays were performed as previous described in (93) with minor changes. Cells were plated in 24 well plates at 3 × 10^3^ cells/well and allowed to settle overnight. Done in triplicate per sample, the corresponding amounts of drugs were added and infected with rAAV 2.5 in equal titers (9.9 × 10^6^ copies/well), then incubated for 7 days. On the 7^th^ day, cells were washed with 1X PBS, fixed with 10% methanol and 10% acetic acid in water for 10-15 min, and stained with 0.1% crystal violet in methanol for 5 min. Plates were then washed with water, allowed to dry overnight, and the crystal violet was resolubilized with 300uL 0.1% SDS in methanol for 2 hrs. 100uL of the resolubilized dye was added to a 96 well, round bottom plate (Griener) and the absorbance was measured using the Gen5 (Biotek) plate reader at 595nm wavelength.

### I-Sce1 Repair Assays

I-Sce1 repair assays were performed as previously described (94) with some minor changes. Cells were plated in 6 well plates 1.75 x 10^6 cells/ well and allowed to settle overnight. Cells were transfected with 1.5ug pBASce-1 and 0.5ug of corresponding pscAAV using Lipofectamine 3000 (Invitrogen) in antibiotic and serum free media. Prior to transfection, cell media was changed to DMEM high glucose (Gibco) and L-glutamine (Gibco) and 5% fetal bovine serum (Gibco) without antibiotics. Cells were allowed to incubate with transfection reaction for 30-48 hrs, then harvested, fixed with 4% PFA, and resuspended in FACS buffer (3% BSA in PBS). At least 20,000 cells/ condition were measured through flow cytometry (Attune NxT) and data was analyzed using FlowJo.

## Acknowledgements

We thank Jeremy Stark for I-SceI reporter cell lines. We thank Koki Morizono for technical assistance. We thank Peter Bradley and Steven Bensinger for use of equipment. We thank Michael Emerman and Toshi Taniguchi for comments on the manuscript.

