## Supplemental Data and Legends for "HIV Vpr modulates the host DNA damage response at two independent steps to damage DNA and repress double-strand DNA break repair"

**Running Title:** HIV activates and represses DNA damage response

**Authors:** Donna Li[a], Andrew Lopez[a,b], Carina Sandoval[a,b], Randilea Nichols Doyle[a], Oliver I Fregoso [a,b]

D.L. and A.L. contributed equally to this work. Author order was determined alphabetically.

[a]Department of Microbiology, Immunology, and Molecular Genetics, University of California, Los Angeles, California, USA.

[b]Molecular Biology Institute, University of California, Los Angeles, California, USA.

### SUPPLEMENTAL LEGENDS & FIGURES

**Figure S1. HIV-1 and HIV-2 Vpr cause cell cycle arrest and expression does not correlate with extent of DNA damage in U2OS cells (data are related to Fig. 1).**

A. Representative bivariate cell cycle analysis flow cytometry plots for propidium iodide (PI, total DNA content) and EdU (DNA synthesis). Gating shows G1, S, and G2 populations of 10,000 U2OS cells treated as in Fig 1A (n=4, one representative experiment shown).

B-C. Mean fluorescence intensities (MFI) of individual cells for DAPI and indicated DNA damage response marker plotted against MFI of 3X-FLAG Vpr. 15-50 cells were measured per condition. Simple linear regression lines are shown. AU = arbitrary units

A. R squared values from linear regression lines in B-C.

**Figure S2. Cell cycle arrest and activation of DNA damage response are conserved by diverse HIV-1 and HIV-2 Vpr isolates (data are related to Fig. 1).**

A. Representative univariate cell cycle analysis flow cytometry plots for uninfected control, empty vector, HIV-1 Q23-17, M.G SE6165 (a group M subtype G sequence), N consensus, O consensus, P consensus, and HIV-2 ROD9, 7312A (a group B sequence), A.PT (a group A sequence), and G.CI.92 (a divergent sequence) using PI staining. Gating shows G1 and G2 populations of 10,000 U2OS cells treated in similar conditions as Fig. 1A (n=2, one representative experiment shown).

B. Box plot representation of the $\gamma$H2AX MFI distribution for uninfected control, empty vector, HIV-1 Q23-17, M.G SE6165, N consensus, O consensus, P consensus, and HIV-2 ROD9, 7312A, A.PT, and GCI.92. 100 cells were measured per condition and treated in conditions similar to Fig. 1A. Asterisks indicates statistical significance from empty vector control as described in Fig. 1B (n=2, one representative experiment shown). AU = arbitrary units.

**Figure S3. DNA replication stalling does not correlate with levels of Vpr expression (data are related to Fig. 2).**

A. Representative DNA replication analysis flow cytometry plots for EdU (DNA synthesis) and Hoechst (total DNA content). Gating shows S phase populations of 10,000 U2OS cells treated in the same conditions as in Fig 1A (n=3, one representative experiment shown).

B. Contour plots of U2OS cells from S phase gate in A, plotted for mCherry and EdU to assay correlation between DNA replication and Vpr expression.

C. Mean Fluorescence Intensities (MFI) for mCherry and EdU from S phase gates in A.

**Figure S4. ATR inhibition, but not ATM inhibition, blocks Vpr-mediated cell cycle arrest (related to Fig. 3).**

A. Representative univariate cell cycle analysis flow cytometry plots for uninfected control, empty vector, HIV-1, HIV-2, and etoposide with or without the ATR inhibitor VE-821 (ATRi) in conditions similar to Fig. 3A, treated for 38 hrs and stained with PI. Gating shows G1 and G2 populations for 10,000 U2OS cells.
B. Representative univariate cell cycle analysis flow cytometry plots for uninfected control, empty vector, HIV-1, HIV-2, and etoposide with the ATM inhibitor KU-55933 (ATMi), as in A. DMSO was used as a control for ATRi and ATMi.

**Figure S5. HIV-1 and HIV-2 Vpr do not alter other DNA repair pathways (related to Fig. 5).**

A. I-SceI SSA assay representing average percent repair by single-strand annealing from four experiments (n=4), normalized to the I-SceI only condition. Cells were treated and analyzed as described in Fig. 5B. Error bars represent +/- standard deviation.
B. I-SceI alt-NHEJ assay representing average percent repair by alternative non-homologous end joining from four experiments (n=4), normalized to the I-SceI only condition. Cells were treated and analyzed as described in Fig. 5B. Error bars represent +/- standard deviation.

**Figure S6. HIV-1 and HIV-2 Vpr mutant analysis (related to Fig. 6).**

A. Mutant Vpr protein expression. Top panel, anti-3X FLAG Vpr; bottom panel, anti-actin as a control for equal total protein loading.
B. Representative bivariate cell cycle analysis flow cytometry plots for propidium iodide (PI, total DNA content) and EdU (DNA synthesis). Gating shows G1, S, and G2 populations of 10,000 U2OS cells treated in the same conditions as in Fig 1A (n=3, one representative experiment shown), corresponds to Fig. 6A.
C. Distribution of the % tail DNA measured for 100 cells per condition from experiment shown in Fig. 6B using the *OpenComet* plug-in. Cells were treated and analyzed as described in Fig. 2B and Fig. 6B.

**Figure S7. HIV-1 and HIV-2 Vpr DCAF1-binding mutant analysis (related to Fig. 7).**

Representative univariate cell cycle analysis flow cytometry plots for uninfected control, empty vector, HIV-1, HIV-2, and mutants. Cells were treated for 38 hrs and stained with PI. Gating shows G1 and G2 populations for 10,000 U2OS cells.

Figure S1

A

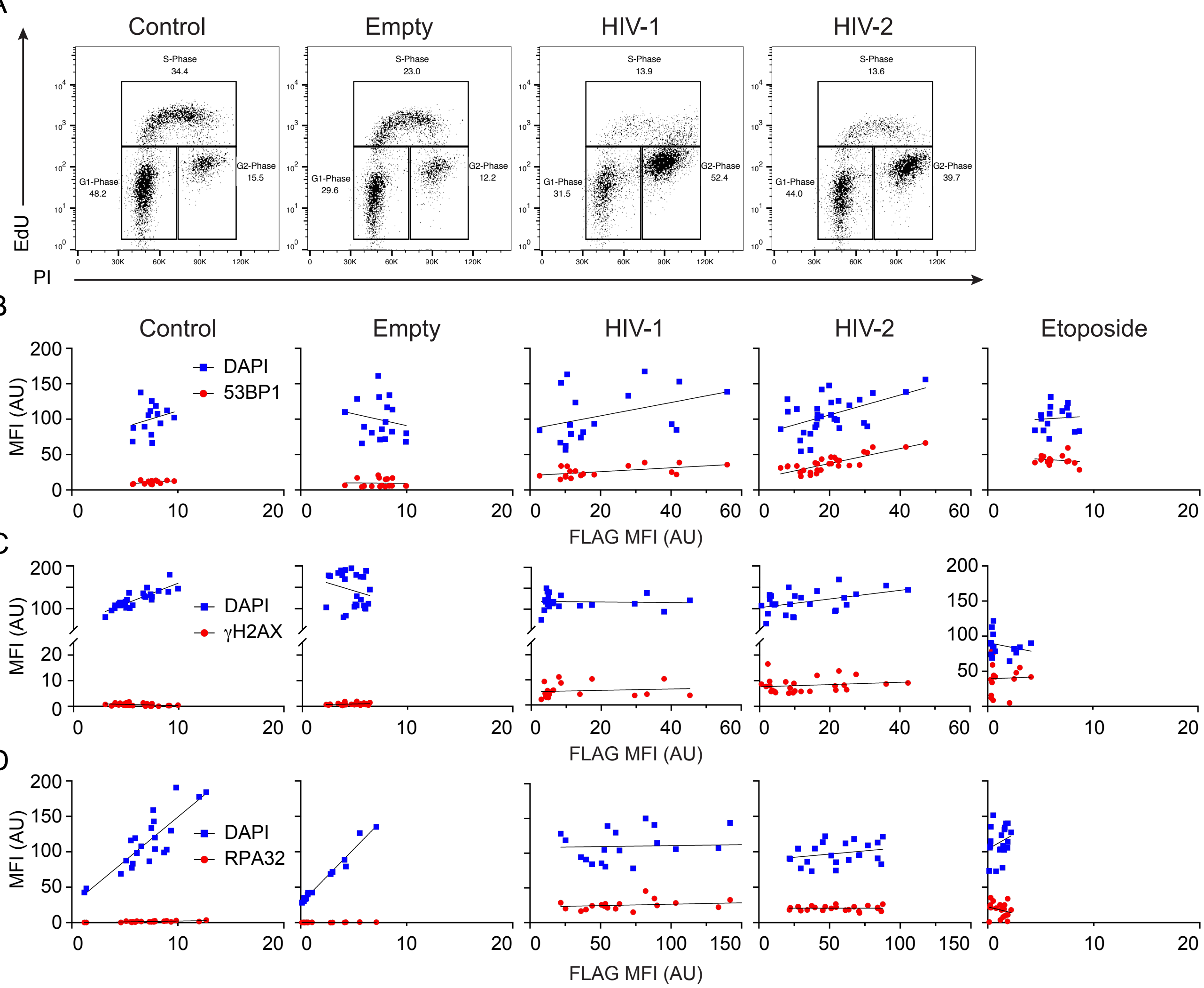

E

### R squared values

| | Control | Empty | HIV-1 | HIV-2 | Etoposide |
|---|---|---|---|---|---|
| **DAPI** | 0.0569 | 0.0341 | 0.5555 | 0.4399 | 0.0047 |
| **53BP1** | 0.1889 | 0.0096 | 0.1071 | 0.1397 | 0.0272 |
| **DAPI** | 0.7261 | 0.0517 | 0.0030 | 0.2006 | 0.0578 |
| **gH2AX** | 0.1442 | 0.0497 | 0.0163 | 0.0296 | 0.0012 |
| **DAPI** | 0.7294 | 0.9776 | 0.0042 | 0.0689 | 0.06499 |
| **RPA32** | 0.5330 | 0.2575 | 0.0891 | 1.257e-006 | 0.06019 |

Figure S2

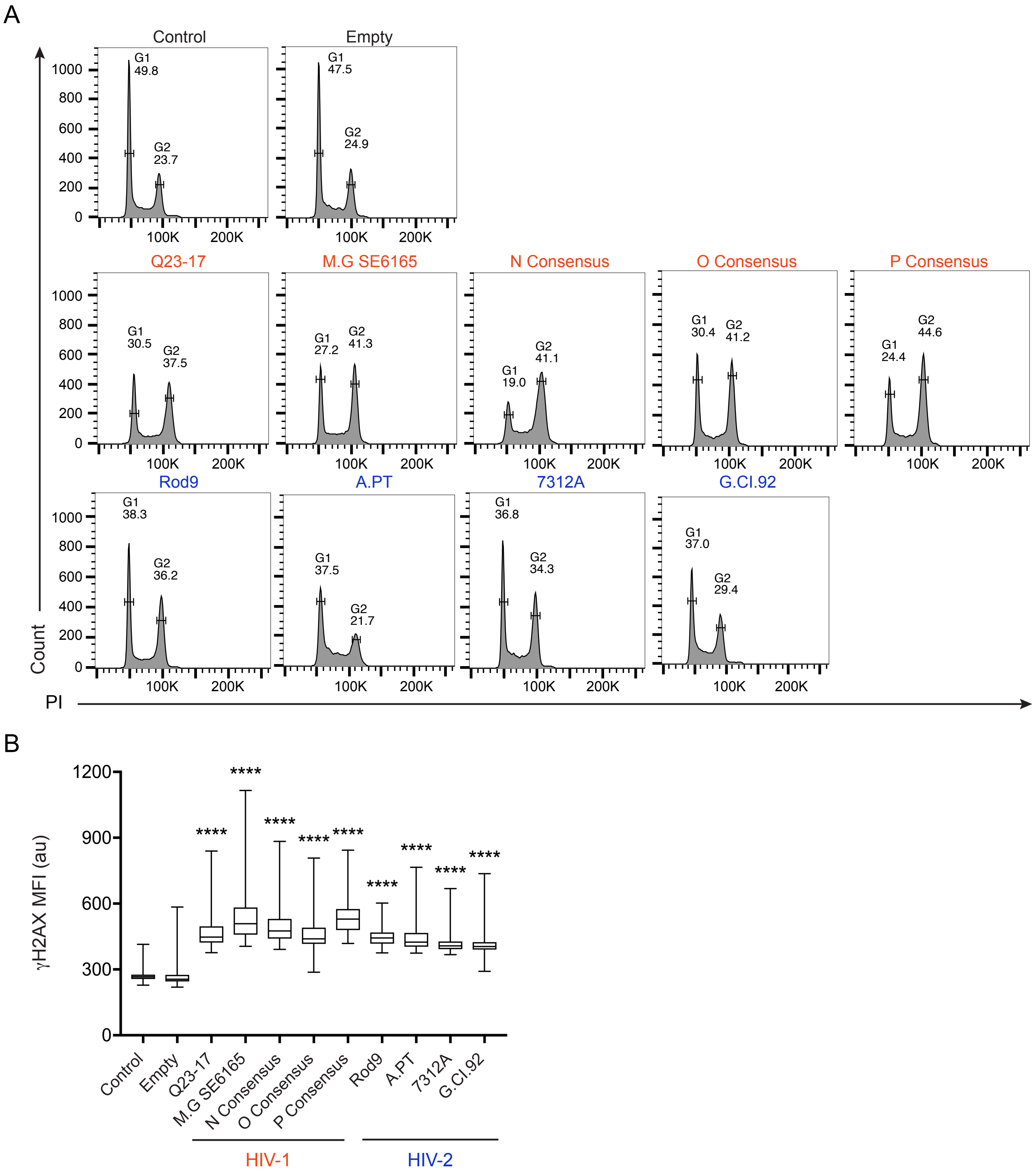
A
Control
Empty
Q23-17
M.G SE6165
N Consensus
O Consensus
P Consensus
Rod9
A.PT
7312A
G.CI.92
G1 49.8
G2 23.7
G1 47.5
G2 24.9
G1 30.5
G2 37.5
G1 27.2
G2 41.3
G1 19.0
G2 41.1
G1 30.4
G2 41.2
G1 24.4
G2 44.6
G1 38.3
G2 36.2
G1 37.5
G2 21.7
G1 36.8
G2 34.3
G1 37.0
G2 29.4
Count
PI
100K
200K
B
γH2AX MFI (au)
1200
900
600
300
0
****
HIV-1
HIV-2

Figure S3

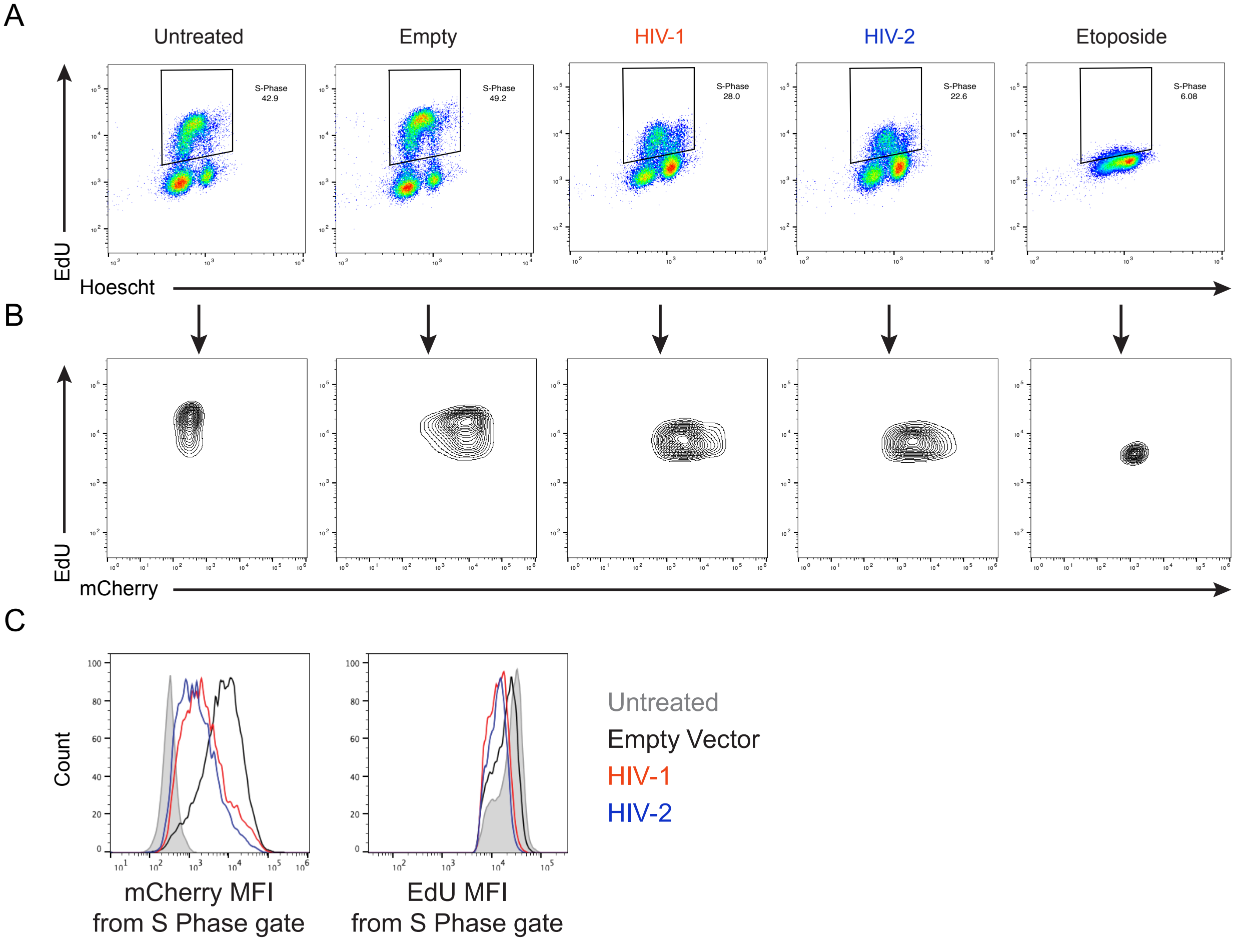
A
Untreated
Empty
HIV-1
HIV-2
Etoposide
S-Phase 42.9
S-Phase 49.2
S-Phase 28.0
S-Phase 22.6
S-Phase 6.08
EdU
Hoescht
B
EdU
mCherry
C
Count
mCherry MFI from S Phase gate
EdU MFI from S Phase gate
Untreated
Empty Vector
HIV-1
HIV-2

Figure S4

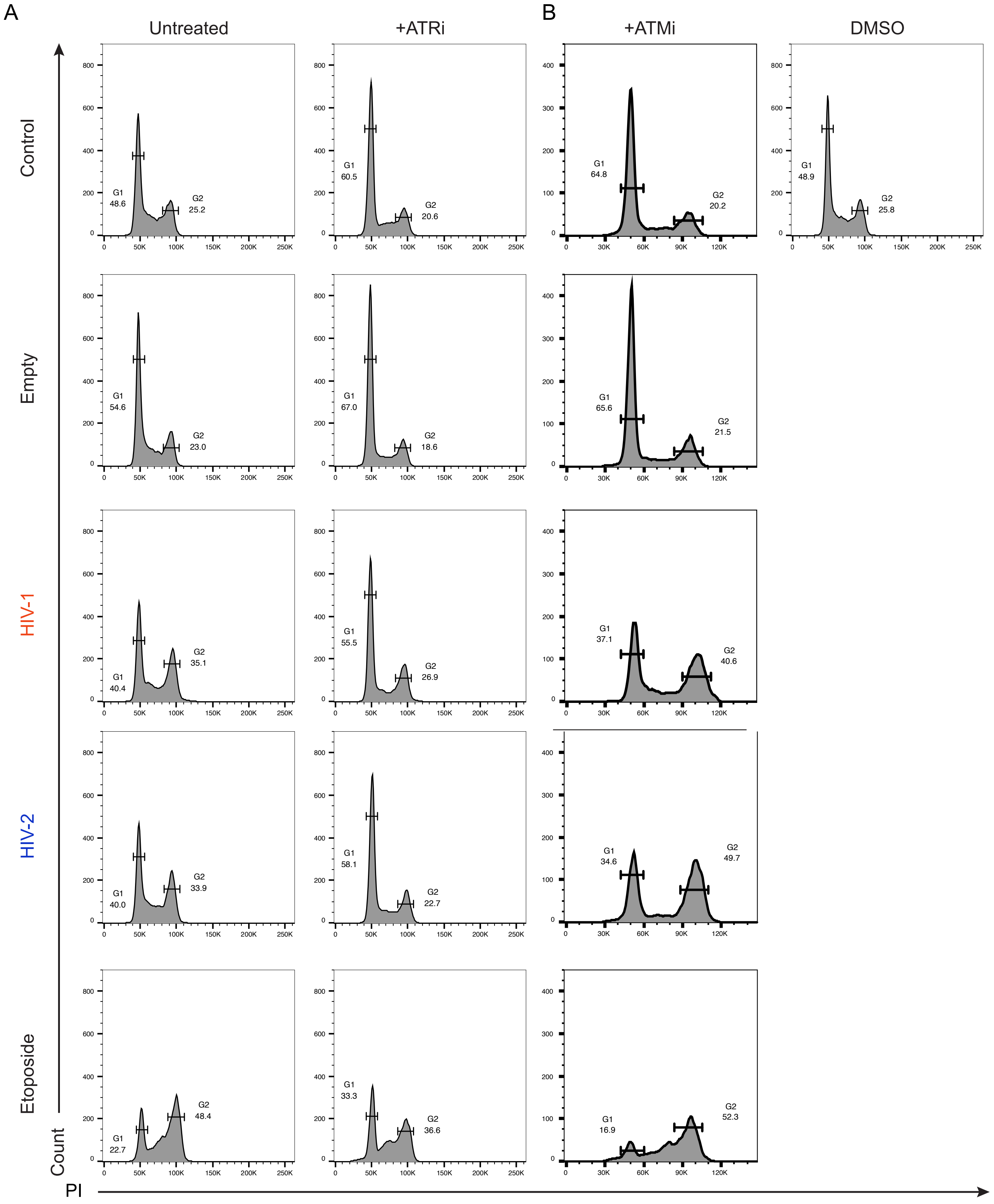
A
B
Untreated
+ATRi
+ATMi
DMSO
Control
Empty
HIV-1
HIV-2
Etoposide
Count
PI
G1 48.6
G2 25.2
G1 60.5
G2 20.6
G1 64.8
G2 20.2
G1 48.9
G2 25.8
G1 54.6
G2 23.0
G1 67.0
G2 18.6
G1 65.6
G2 21.5
G1 40.4
G2 35.1
G1 55.5
G2 26.9
G1 37.1
G2 40.6
G1 40.0
G2 33.9
G1 58.1
G2 22.7
G1 34.6
G2 49.7
G1 22.7
G2 48.4
G1 33.3
G2 36.6
G1 16.9
G2 52.3

Figure S5

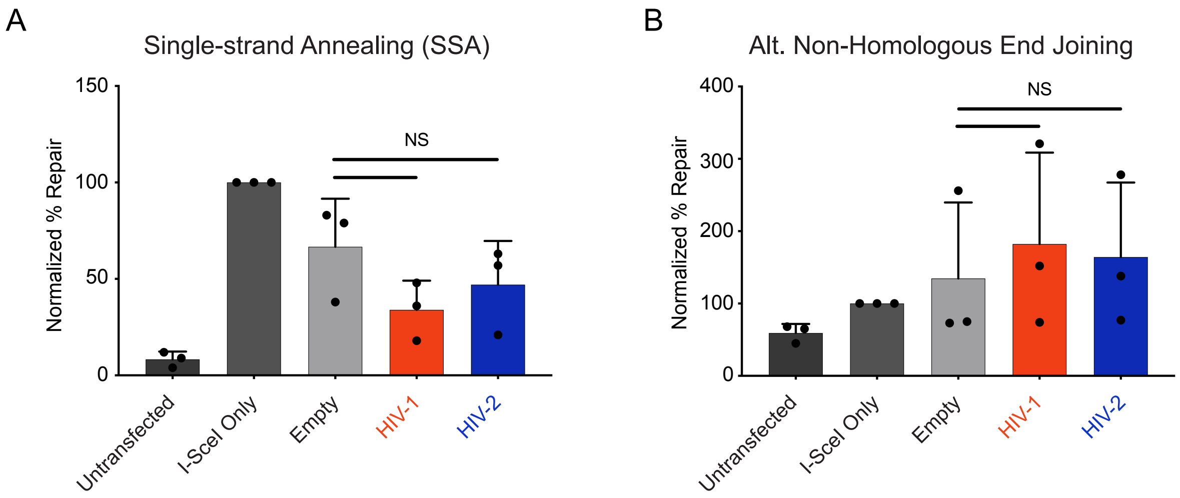
A
Single-strand Annealing (SSA)
Normalized % Repair
150
100
50
0
NS
Untransfected
I-SceI Only
Empty
HIV-1
HIV-2
B
Alt. Non-Homologous End Joining
Normalized % Repair
400
300
200
100
0
NS
Untransfected
I-SceI Only
Empty
HIV-1
HIV-2

Figure S6

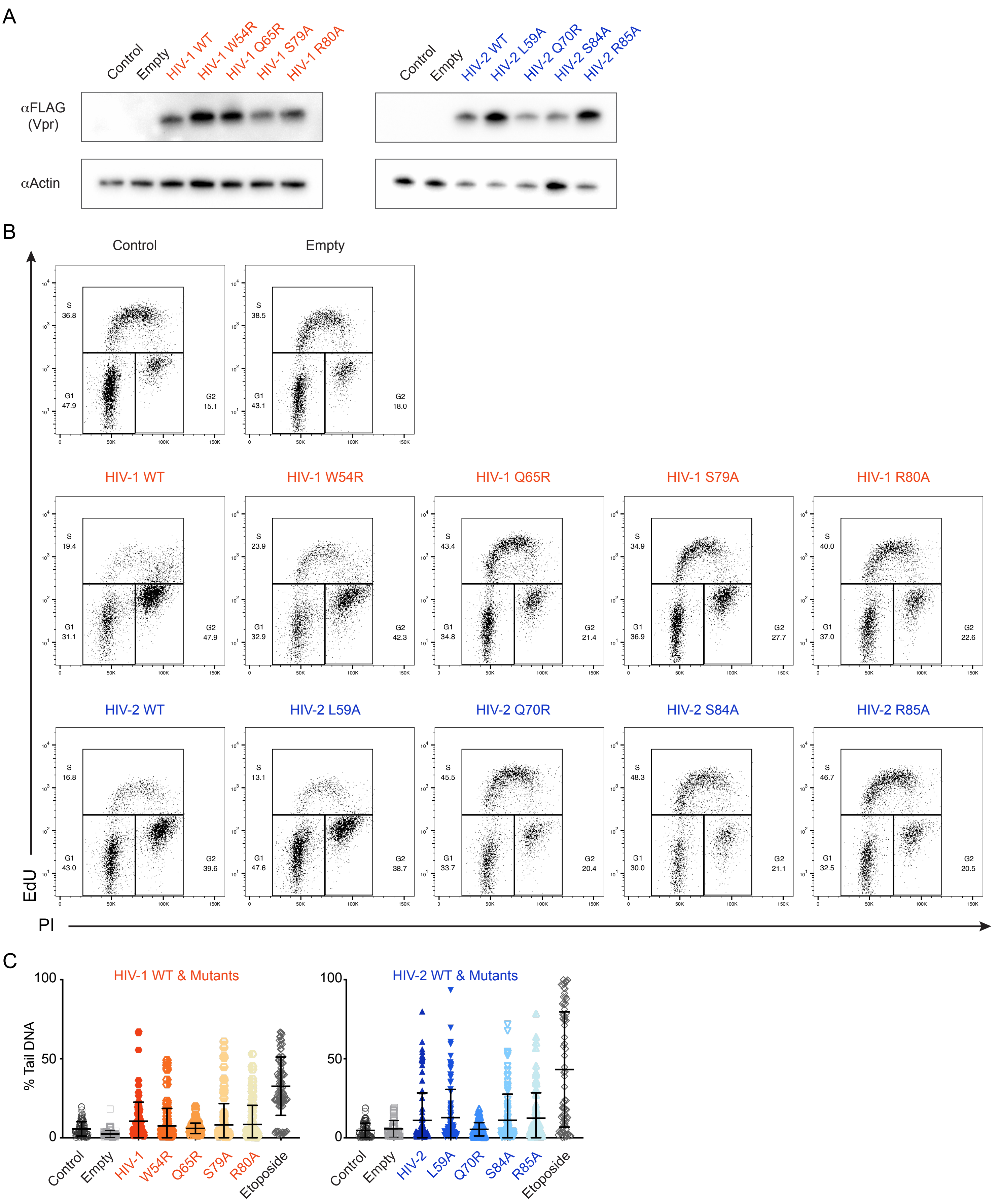
A
Control
Empty
HIV-1 WT
HIV-1 W54R
HIV-1 Q65R
HIV-1 S79A
HIV-1 R80A
HIV-2 WT
HIV-2 L59A
HIV-2 Q70R
HIV-2 S84A
HIV-2 R85A
αFLAG (Vpr)
αActin
B
EdU
PI
C
HIV-1 WT & Mutants
HIV-2 WT & Mutants
% Tail DNA
Etoposide

Figure S7

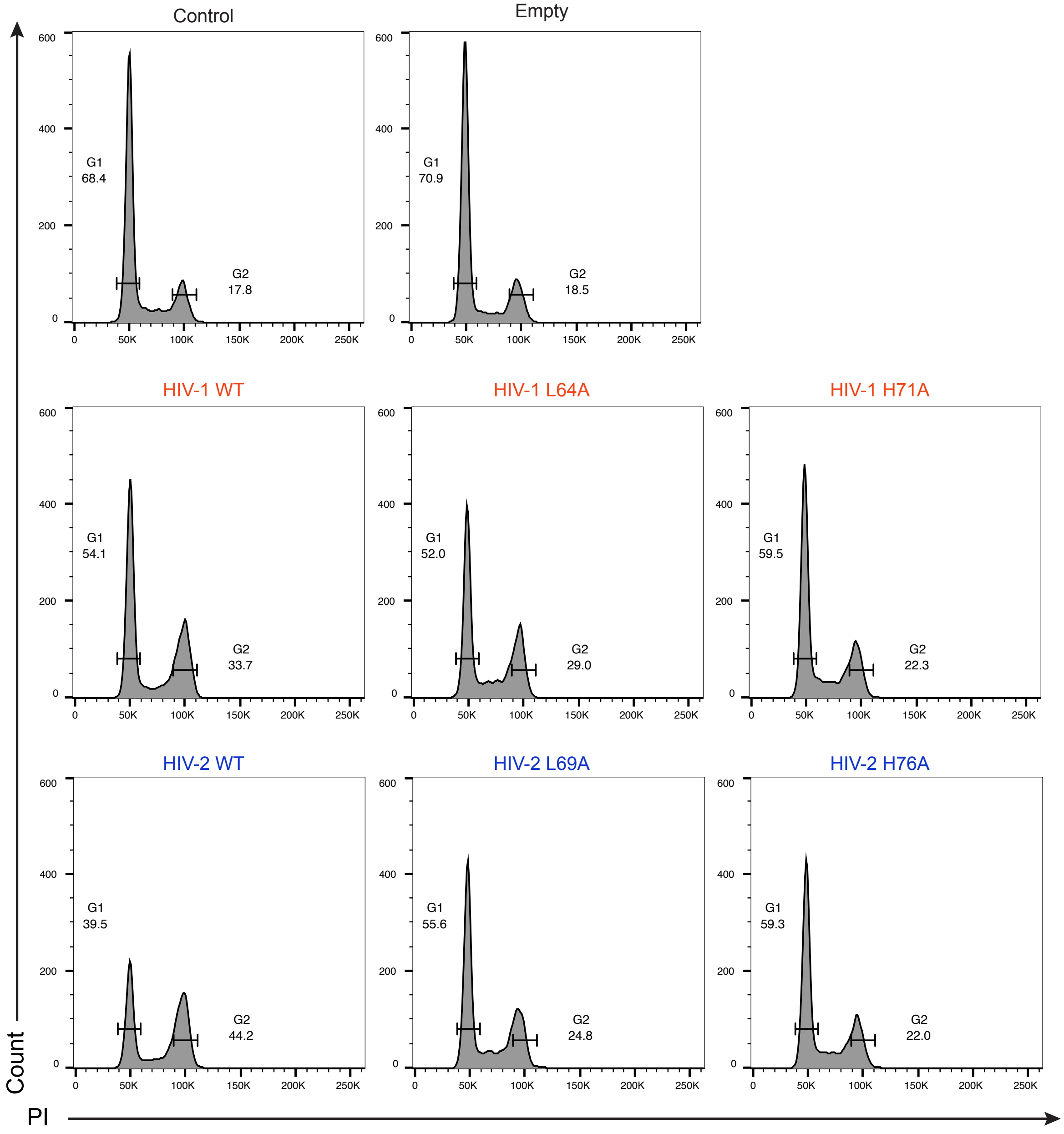

Control
Empty
HIV-1 WT
HIV-1 L64A
HIV-1 H71A
HIV-2 WT
HIV-2 L69A
HIV-2 H76A
G1
68.4
G2
17.8
G1
70.9
G2
18.5
G1
54.1
G2
33.7
G1
52.0
G2
29.0
G1
59.5
G2
22.3
G1
39.5
G2
44.2
G1
55.6
G2
24.8
G1
59.3
G2
22.0
Count
PI
